# High-throughput inverse toeprinting reveals the complete sequence dependence of ribosome-targeting antibiotics

**DOI:** 10.1101/298794

**Authors:** Britta Seip, Guénaёl Sacheau, Denis Dupuy, C. Axel Innis

## Abstract

It has recently become clear that various antibiotics block the translation of bacterial proteins in a sequence-specific manner. In order to understand how this specificity contributes to antibiotic potency and select better antimicrobial leads, new high-throughput tools are needed. Here, we present inverse toeprinting, a new method to map the position of ribosomes arrested on messenger RNAs during *in vitro* translation. Unlike ribosome profiling, our method protects the entire coding region upstream of a stalled ribosome, making it possible to work with transcript libraries that randomly sample the sequence space. We used inverse toeprinting to characterize the pausing landscape of free and drug-bound bacterial ribosomes engaged in translation. We obtained a comprehensive list of arrest motifs that could be validated *in vivo*, along with a quantitative measure of their pause strength. Thus, our method provides a highly parallel and scalable means to characterize the sequence specificity of translation inhibitors.

---

Ribosome-targeting antibiotics, such as macrolides, phenicols or oxazolidinones, arrest translation in a context-dependent manner that can be attributed to specific amino acid motifs within the nascent polypeptide^1–6^. Although the link between the sequence specificity of these antibiotics and their efficacy as ribosomal inhibitors remains to be established, a better understanding of this process could help pinpoint the most promising lead compounds for further development by accelerating their characterization. In order to do so, however, high-throughput tools that can reveal the complete sequence-dependence of ribosomal inhibitors are needed. This is particularly true for macrolide antibiotics, which have seen a resurgence following the introduction of a total synthesis approach that allows the production of a virtually endless number of derivatives^7, 8^.

At present, ribosome profiling^9^ is the only tool capable of providing a global *in vivo* view of ribosome density on the mRNA, which can be used as a proxy to measure context-dependent translation inhibition. Ribosome profiling has been used to study the translational pausing landscape in EF-P-deficient bacteria^10^ and to identify nascent amino acid motifs responsible for antibiotic-dependent translational arrest in the Gram-negative bacterium *Escherichia coli*^1^ and in the Gram-positive bacterium *Staphylococcus aureus*^2^. However, the bacterial culture volumes and sequencing depth necessary to perform ribosome profiling are incompatible with the systematic screening of potential antibiotic leads. To overcome this limitation, we developed a new, highly scalable *in vitro* method to systematically characterize translational arrest by ribosome-targeting compounds.

Inverse toeprinting is a versatile selection strategy that relies on a highly processive 3’ to 5’ RNA exonuclease to degrade the mRNA downstream of the leading ribosome on a transcript. This makes it possible to determine the position of stalled ribosomes on the mRNA with codon resolution, while protecting the entire upstream peptide-encoding region. Inverse toeprinting is amenable to high-throughput, as next-generation sequencing can be used as readout. Since it is an *in vitro* method, it allows the precise control of translation conditions and can further be incorporated into selection schemes tailored to a variety of applications. We used inverse toeprinting to explore the pausing landscape of free and drug-bound bacterial ribosomes engaged in the translation of a library of random transcripts, and demonstrate its usefulness for rapidly assaying context-specific translation inhibition by ribosome-targeting compounds.

## Results

### Inverse toeprinting maps stalled ribosomes with codon resolution while preserving the upstream coding region

Ribosome profiling only generates a short footprint and thus loses sequence information for most of the coding region, while classical toeprinting^11, 12^ can accurately determine the position of the ribosome on the mRNA, but is not suitable for large-scale analyses (**Fig. 1a**). In contrast, inverse toeprinting relies on ribonuclease R, a highly processive 3’ to 5’ RNA exonuclease that efficiently degrades mRNA featuring a 3’ poly–(A) tail^13^. Our assumption was that RNase R would digest the mRNA up to a precise, discrete position downstream of the P-site codon of the foremost stalled ribosome-nascent chain complex. Inverse toeprints obtained in this manner could be analyzed by deep sequencing or evolved through multiple rounds of selection (**Fig. 1b, Supplementary Fig. 1a**).

**Fig. 1.**
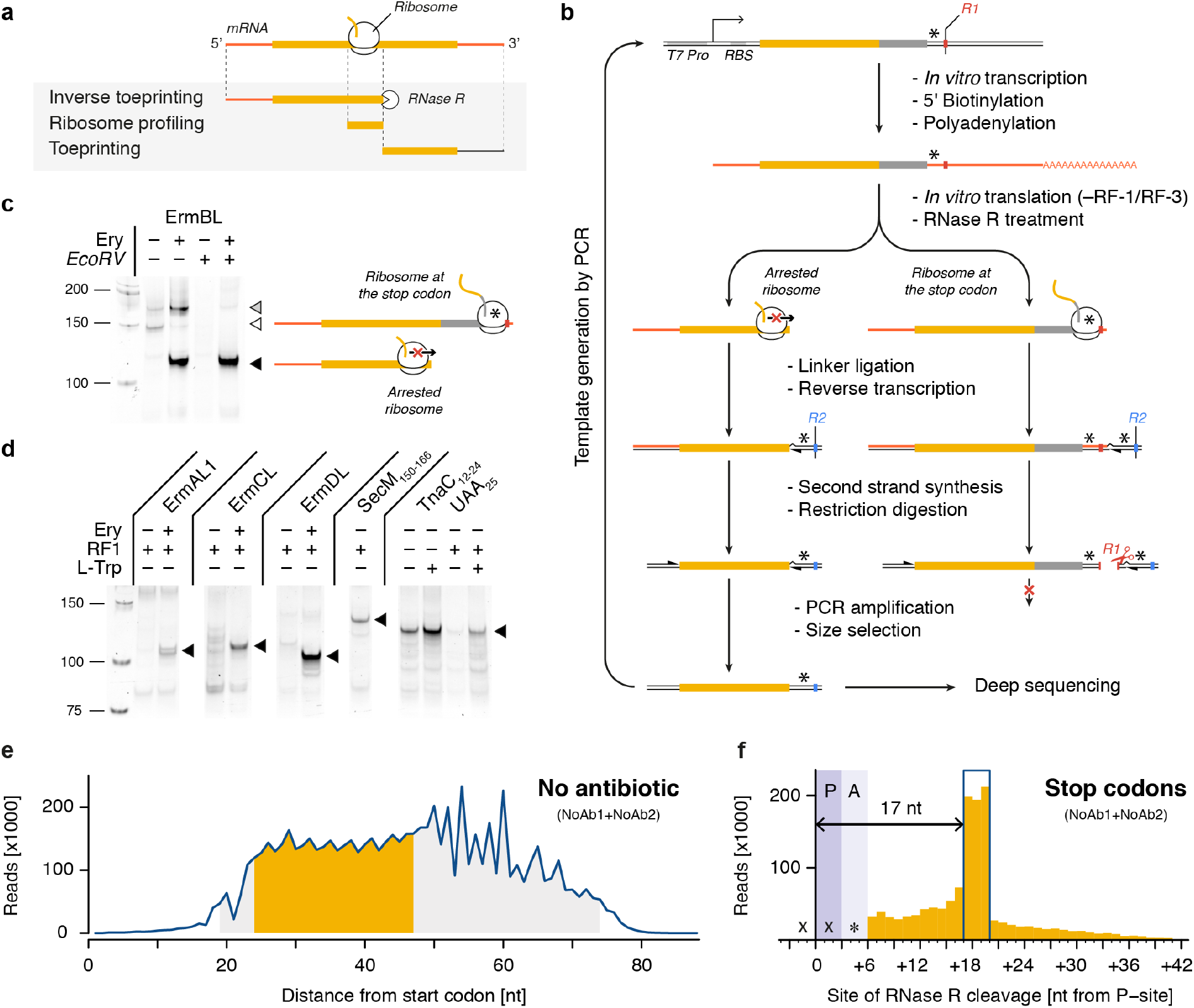
Inverse toeprinting locates ribosomes on the mRNA with codon resolution. (a) Comparison between inverse toeprinting, ribosome profiling and classical toeprinting. (b) Schematic overview of the inverse toeprinting workflow. Restriction enzymes used in odd (*EcoRV*) and even (*ApoI*) cycles are shown in red and blue, respectively. Stop codons are indicated as asterisks. (c) Removal of inverse toeprints featuring ribosomes that have reached the stop codon on the *ermBL* template (white triangle) using the *EcoRV* restriction enzyme. The black triangle corresponds to arrested ribosomes and the gray triangle to full-length mRNA. (d) Inverse toeprints for various Erm peptides, SecM_150-166_ and TnaC_12-24_UAA_25_. The wild-type UGA stop codon for TnaC was replaced with a UAA stop codon, allowing its release by RF-1. (e) Size distribution of inverse toeprints from two biological replicates (NoAb1 and NoAb2) with a minimum Q-score of 30 obtained from an NNS_15_ library translated in the absence of any added antibiotic. The fragment size range shaded in gray corresponds to the band that was cut from a 12% TBE-acrylamide gel, while the range in yellow indicates fragments that were used in the subsequent analysis. (f) Analysis of inverse toeprints containing stop codons that were obtained in the absence of antibiotic reveals that RNase R cleaves +1 7 nucleotides downstream from the P-site.

As a proof of principle for inverse toeprinting, we generated a 5’–biotinylated (**Supplementary Fig. 1b**) and 3’–polyadenylated (**Supplementary Fig. 1c**) mRNA template encoding the Erythromycin Resistance Methyltransferase B leader peptide (ErmBL), followed by a short coding region ending with a UGA stop codon. We then used this template to drive the expression of ErmBL in a PURE *E. coli* translation system^14^ in the absence of release factor 2 (RF-2), whose physiological role is to promote peptide release at UGA or UAA stop codons^15^. ErmBL belongs to the Erm family of arrest peptides, a group of nascent polypeptides which force the ribosomes that are translating them to stall on the mRNA in an antibiotic-dependent manner^5, 6^. This, in turn, induces the expression of a methyltransferase gene on the same mRNA that ultimately confers resistance to macrolide antibiotics^16^. We chose ErmBL because it has been extensively characterized biochemically^17, 18^ and its structure has been determined within the context of a drug-stalled ribosome^6, 19^. In the presence of the macrolide antibiotic erythromycin (Ery), ribosomes undergo strong translational arrest when codon 10 of *ermBL* is in the ribosomal P-site, whereas ribosomes translating the same sequence in the absence of the drug are able to reach the UGA stop codon. Treatment of both samples with RNase R resulted in short or long 3’–truncated mRNA fragments for ribosomes arrested at these two positions, respectively (**Fig. 1c, Supplementary Fig. 1c**). Ribosome-protected mRNAs were ligated to an oligonucleotide linker through their 3’ ends to enable reverse transcriptase priming. Ribosomes that reached the UGA stop codon protected an *EcoRV* site on the mRNA, which could be cleaved after reverse transcription and second strand synthesis. As a result, only short cDNA fragments derived from messengers encoding sequences that caused drug-dependent translational arrest were amplified by PCR after the *EcoRV* treatment (**Fig. 1c**). Similar experiments performed with other arrest peptides (ErmAL1, ErmCL, ErmDL, SecM and TnaC) showed that inverse toeprinting can be used as a general tool for selecting arrest sequences (**Fig. 1d**). Moreover, we showed that it is possible to perform consecutive rounds of selection by alternating restriction enzymes on the 3’ oligonucleotide linkers (**Supplementary Fig. 1d**). This could be used in the future as the basis for a SELEX^20, 21^-like scheme to identify rare arrest sequences contained within a complex transcript pool.

To test the scalability of our method for high-throughput screens, we performed inverse toeprinting on an *in vitro* translation reaction using a template library (NNS_15_ library) encoding 20-residue peptides with a variable region of 15 NNS codons (**Supplementary Fig. 2a**). We size-selected inverse toeprints so as to minimize contamination from DNA fragments resulting from initiation complexes, which appeared as a prominent band on a TBE-polyacrylamide gel (**Supplementary Fig. 3**). Paired-end Illumina sequencing revealed a tri-nucleotide periodicity for fragments where RNase R cleavage had occurred 24 to 47 nucleotides downstream of the start codon (**Fig. 1e**). Longer fragments did not follow this size periodicity and were excluded from our analysis.

Despite the presence of release factor 1 (RF-1) in the translation reaction, we noticed that UAG stop codons recognized by this termination factor^15^ were enriched 1.4-fold upon selection, indicating that ribosomes linger on these codons. We used this observation to precisely characterize the protective effect of stalled ribosomes on the mRNA. Given that ribosomes stall when they encounter a UAG stop in the A-site, our analysis revealed a distinct three-nucleotide peak positioned +17 nucleotides downstream from the P-site (**Fig. 1f**). This RNase R cleavage pattern could also be confirmed for all of the known arrest motifs we tested, even allowing us to locate a second stalling site for the drug-dependent arrest peptide ErmAL1 that had been overlooked in previous studies^5, 22^ (**Supplementary Fig. 4a-f**). This demonstrates that inverse toeprinting can be used to determine the position of stalled ribosomes on the mRNA at codon resolution.

### Motif enrichment correlates with pause strength and *in vivo* data

In order to assess whether the enrichment of sequences following inverse toeprinting is correlated to their ability to induce translational arrest or pausing, we first sought to obtain a subset of well-measured 3-amino acid (3-aa) motifs. To do so, we estimated the reproducibility of the frequency of occurrence of 3-aa motifs between independent biological replicates. From ~3.4 million inverse toeprints obtained after translation in the absence of antibiotic, we could precisely (Supplementary Fig. 5) and reproducibly measure the frequency of 5,278 of 8,000 possible 3-aa motifs (*R^2^*=0.95; <15% error between biological replicates) (**Fig. 2a**). In order to limit the impact of the noise resulting from poor counting statistics, we chose to limit our subsequent analysis to this subset of well-measured 3-aa motifs.

**Fig. 2.**
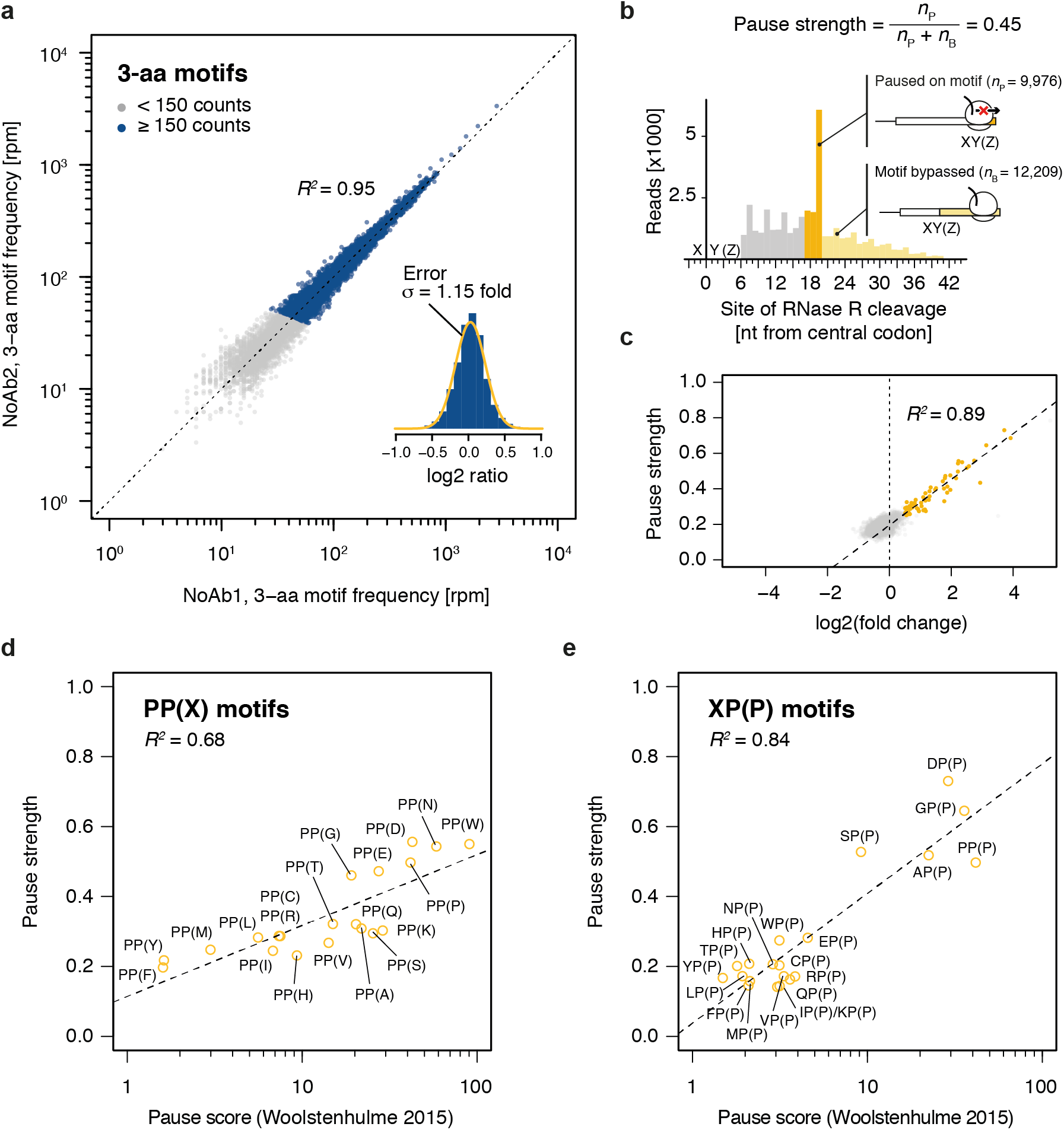
Motif pause strength correlates with enrichment upon inverse toeprinting. (a) 3-aa motif frequencies in reads per million (rpm) from two independent inverse toeprinting experiments performed after translation in the absence of antibiotic (NoAb1 and NoAb2). The inset represents a histogram of log2 ratios between replicates for 3-aa motifs having low statistical counting error (i.e. with >150 counts (blue), Supplementary Fig. 5), with an overlaid normal error curve (mean = 0.02, standard deviation = 0.2 log2 units, equivalent to σ= 1.15 fold). (b) Formula used to calculate pause strength for an XY(Z) motif, with the amino acid in the ribosomal A-site in brackets. (c) Plot of pause strength against log2(fold change) of all possible 3-aa motif frequencies relative to the NNS_15_ library. Yellow points correspond to intrinsic 3-aa pause motifs with a pause strength ≥ 0.25. All other motifs are shown as gray dots. (d, e) Plot of pause strengths calculated in this study against pause scores calculated from ribosome profiling data obtained from *E. coli* cells lacking EF-P, for (d) PP(X) and (e) XP(P) motifs^10^. The scores obtained by both methods are strongly correlated, as indicated by *R^2^* values of 0.68 and 0.84 for PP(X) and XP(P) motifs, respectively.

Next, we devised a means to quantify the strength of translational pausing for each individual 3-aa motif, which we call the pause strength (**Fig. 2b**). We calculated pause strengths for all reliably measured 3-aa motifs and observed that pause strength showed a strong linear correlation with the log2(fold change) in frequency for all motifs exhibiting a pause strength greater than 0.25 (**Fig. 2c**). Pause strengths were in the range of 0.1–0.8, with values below 0.25 found for a majority of motifs. In the absence of antibiotic, PP(X) and XP(P) motifs displayed pause strengths that were strongly correlated with the “pause scores” calculated for the same motifs from ribosome profiling experiments performed in EF-P-deficient *E. coli*^10^ (**Fig. 2d, e**). While PP(X) motifs on the whole appear to be more efficient at pausing translation (PP(D) > PP(W) > PP(N) > PP(P) > PP(E) > PP(G)), the strongest XP(P) motifs (DP(P) > GP(P) > SP(P) > AP(P) >PP(P)) exhibit the greatest pause strengths. Interestingly, we could identify a new intrinsic arrest motif (XP(C)), where X corresponds to the amino acids in XP(P) motifs that cause the strongest pauses (i.e. A, D, G, S). Thus, inverse toeprinting provided us with a detailed view of the intrinsic translational pausing landscape of *E. coli* ribosomes that matches earlier *in vivo* profiling data^10^. Critically, we showed that motif enrichment upon inverse toeprinting and pause strength are strongly correlated, indicating the robustness of the selection procedure.

### Pause-inducing motifs are reproducibly enriched following translation in the presence of erythromycin

We further sought to perform a comprehensive identification of short 3-aa motifs that arrest ribosomes in response to the macrolide antibiotic Ery. Previous studies showed that ribosomes translating in the presence of Ery stall when they encounter +X(+) motifs, where “+” stands for the positively charged amino acids arginine or lysine^1, 2, 23^. We therefore performed inverse toeprinting on an *in vitro* translation reaction using the NNS_15_ library in the presence of Ery. Paired-end Illumina sequencing of the corresponding inverse toeprints revealed a much stronger tri-nucleotide periodicity of fragment sizes compared to the samples without antibiotic (**Supplementary Fig. 6**). Comparison of independent biological replicates of inverse toeprints obtained in the presence of Ery led to the identification of a subset of 3-aa motifs that were well measured and used for subsequent analyses (**Supplementary Fig. 7**).

From this subset, we measured the enrichment of 3-aa motifs in inverse toeprints obtained after translation in the presence of Ery relative to those obtained in the absence of antibiotic, and calculated a mean error of 1.2-fold change in motif frequency upon addition of Ery (**Fig. 3a**). A number of 3-aa motifs that were enriched significantly in the Ery sample were characterized by relatively high pause strengths (**Fig. 3a, c**, **Supplementary Fig. 8**, **Supplementary Fig. 9b**), which was not the case in the absence of the drug (**Fig. 3b**). These include the previously reported +X(+) motif, the general XP(X) motif and its subset XP(W), as well as the +X(W) motif.

**Fig. 3.**
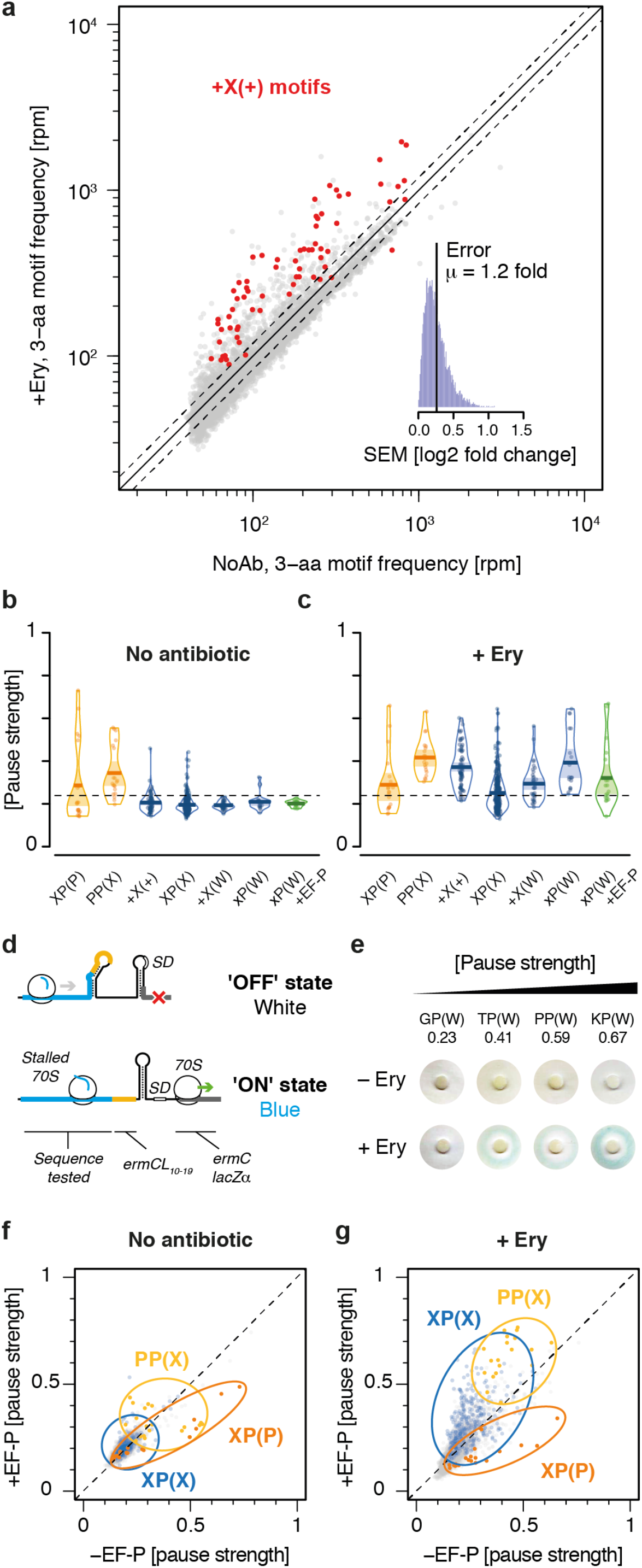
Nascent peptide-dependent translational arrest in response to Ery. (a) Frequency of occurrence of 3-aa motifs with low statistical counting error in inverse toeprints obtained in the absence or presence of Ery, with +X(+) motifs indicated in red. The inset represents a histogram of the standard error of the mean (SEM) of the log2 fold change in 3-aa motif frequency upon addition of Ery. The upper and lower dotted lines (gray) indicate 1.20 and 0.83-fold changes, respectively, corresponding to the mean (μ) of the distribution of SEM(log2 fold change). (b, c) RDI (Raw data, Description and Inference) plot showing pause strengths for individual motifs translated in the (b) absence or (c) presence of Ery. Polyproline motifs are shown in yellow, xP(W) motifs for the +EF-P sample are in green and all other motifs are in blue. The horizontal dashed line corresponds to the 0.25 pause strength cutoff used to identify motifs that are enriched upon addition of Ery. (d) Overview of the *lacZ*α-complementation assay used to test the *in vivo* activity of ErmBL variants (modified from Bailey *et al*.^26^). (e) Disc-diffusion test plates used to assay the ability of nascent formyl-MAXP(W) to cause translational arrest *in vivo* in an *E. coli* strain expressing EF-P. Discs marked with +Ery contain this antibiotic and blue rings result from the induction of a *lacZ*α reporter in response to ribosome stalling at an upstream test ORF (modified from Bailey *et al*.^26^). (f, g) Plots of pause strength in the presence of EF-P against pause strength in the absence of EF-P, for samples (f) without antibiotic or (g) with Ery. XP(P) motifs are indicated in orange, PP(X) motifs in yellow and XP(X) motifs in blue. All other motifs are in gray.

Ribosomal pausing at XP(W) motifs in the presence of Ery was reported earlier^1^, but we show here that ribosomes stalled on these motifs are not significantly rescued by the translation factor EF-P^24, 25^, as evidenced by inverse toeprinting in the presence of EF-P (**Fig. 3c**) and by stalling assays in *E. coli* expressing EF-P (**Fig. 3d, e**). Interestingly, a majority of XP(X) motifs induced significant drug-dependent pauses in the presence of EF-P, but not in its absence (**Fig. 3f, g**), suggesting that the effects of the drug are exacerbated in an EF-P competent background. XP(P) and PP(X) motifs also behaved differently in the presence of drug, with pausing at XP(P) motifs being efficiently rescued by EF-P, while PP(X) motifs induced greater pausing when EF-P was added to the reaction (**Fig. 3g**). The molecular basis for this phenomenon is unclear at present and will need to be further investigated.

### Inverse toeprinting identifies ErmBL variants that discriminate between closely related antibiotics

To test the selective power of our method, we performed inverse toeprinting on a high complexity transcript library encoding variants of the macrolide-dependent arrest peptide ErmBL (**Supplementary Fig. 2b**), in the absence or presence of Ery (**Fig. 4a**) or of the weaker antibiotic oleandomycin (Ole) (**Fig. 4b**). After paired-end Illumina sequencing, ~1.3 million library reads were aligned to the wild-type *ermBL* sequence, corresponding to 724,573 unique protein variants. Moreover, the observed distribution of mutations within the library closely approximated the expected distribution, with ~91% of single mutants and ~45% of double mutants sequenced. Inverse toeprints obtained after translation in the presence of Ery or Ole featured >230,000 unique protein variants in each case, ~80% of which were sequenced more than once.

**Fig. 4.**
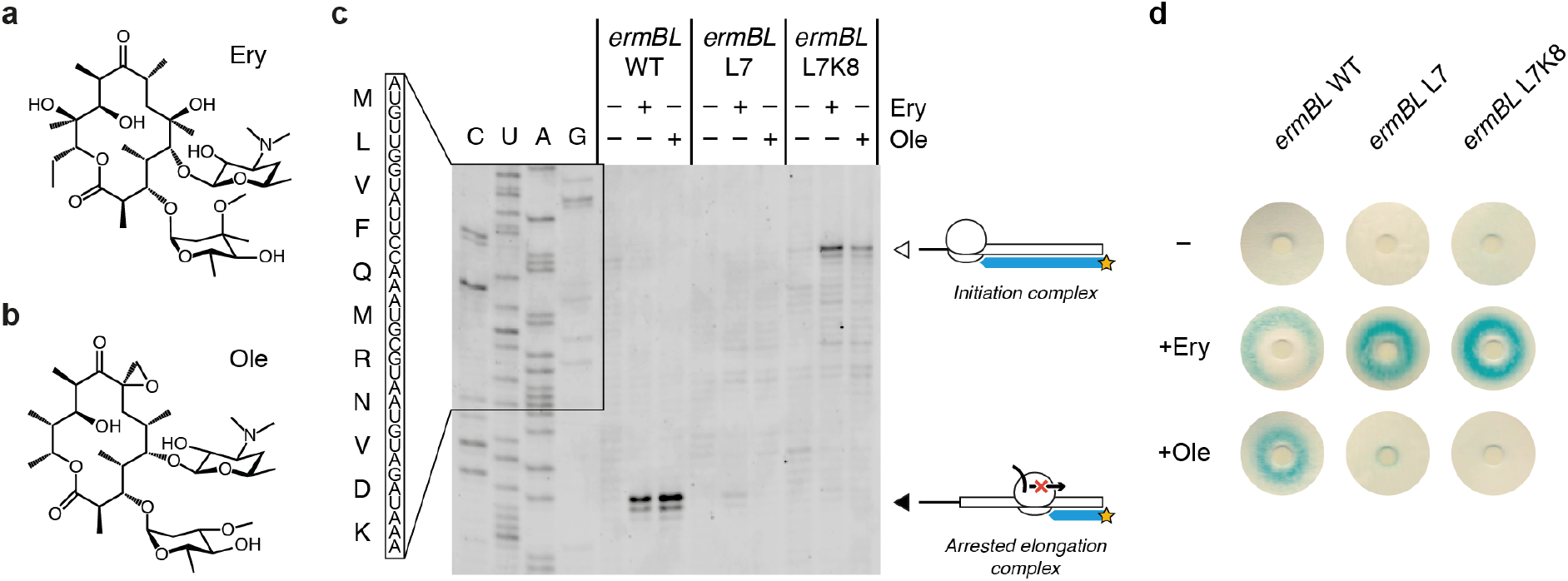
The ErmBL L7 and L7K8 mutants discriminate between closely related antibiotics. Chemical diagrams for (a) Ery and (b) Ole. (c) Classical toeprinting analysis of translational arrest by wild-type ErmBL (*ermBL* WT), an L7 single mutant (*ermBL* L7) and an L7K8 double mutant (*ermBL* L7K8), in the absence or presence of the antibiotics Ery and Ole. The white arrow indicates ribosomes on the start codon and the black arrow indicates arrested elongation complexes with the GAU codon encoding Asp-10 in the ribosomal P-site. The sequence of wild-type ErmBL is shown. (d) Disc-diffusion test plates used to assay the ability of nascent ErmBL WT, ErmBL L7 and ErmBL L7K8 to cause translational arrest *in vivo* in the absence or presence of Ery or Ole soaked into paper discs. It should be noted that the double mutant shows greater antibiotic selectivity *in vivo* compared to the single mutant.

By comparing sequence variants that were obtained after inverse toeprinting in the presence of Ery or Ole, we identified amino acids that were over- or under-represented at each position of ErmBL in response to these antibiotics (**Supplementary Fig. 10**). In doing so, we noticed that leucine is enriched at position 7 of ErmBL (−3 when counting from the amino acid in the P-site) in the presence of Ery, but not Ole. A closer look at the sequences containing leucine at this position revealed several mutants that underwent translational arrest exclusively in the presence of Ery (**Supplementary Table 1**). Among these, we chose to further characterize the L7 single mutant and L7K8 double mutant of ErmBL, both of which were enriched ~ 1.9-fold in the Ery sample compared to the input library whereas they were not significantly enriched in the Ole sample (0.7-fold and 0.5-fold, respectively). Moreover, the L7 and L7K8 mutants originated from 198 and 55 unique variants at the nucleotide level, respectively, indicating multiple independent events. Using an *in vitro* toeprinting assay (**Fig. 4c**) and an *in vivo* reporter to measure translational arrest^26^ (**Fig. 4d**), we confirmed the antibiotic specificity for the L7 and L7K8 mutants. Interestingly, the barely detectable Ery-dependent toeprints obtained *in vitro* contrasted with strong β-galactosidase activity for the *in vivo* assay, indicating that arrest sequences that appear weak by toeprinting can stall ribosomes effectively *in vivo*. Importantly, the L7 and L7K8 mutants could not have been predicted on the basis of the available structure of a stalled ErmBL-70S complex^19^ and would not have been identified using a simple alanine mutagenesis scan of ErmBL. The weak toeprint signals would also likely have been overlooked. This highlights the value of our method for exploring the sequence space of known arrest peptides or for identifying specific variants out of a complex library.

## Discussion

We developed inverse toeprinting, an *in vitro* method that can locate, with codon resolution, ribosomes that are paused or stalled on the mRNA and preserves the entire mRNA sequence upstream of the point of arrest. This presents a major advantage over current methods like ribosome profiling when a reference genome is not available for mapping protected mRNA footprints but the encoded amino acid sequence needs to be known, for example with a random transcript library. Here, we showed that inverse toeprinting is a versatile selection tool that can be used for a variety of applications, including the characterization of the intrinsic translational pausing landscape of the bacterial ribosome, the elucidation of antibiotic-dependent translational arrest profiles or the identification of arrest peptide variants with different drug specificities. Importantly, inverse toeprinting can provide comparable results to *in vivo* methods like ribosome profiling, while providing a greater coverage of sequence space and making it possible to measure the effect of tens if not hundreds of different reaction conditions in a highly parallel fashion. As a result, it provides a convenient and effective means to characterize novel protein synthesis inhibitors and provides a solid basis for developing more complex methods.

Although we have shown that the selection procedure itself is efficient and fully adapted to the study of ribosome-targeting antibiotics, there are a number of key developments to be made for inverse toeprinting to form the basis of a SELEX-like procedure. Such a procedure would be useful to develop novel arrest peptides that can regulate gene expression in response to small molecules. Indeed, naturally occurring arrest peptides like TnaC^27^ in *E. coli* or the fungal arginine attenuator peptide (AAP)^28^ arrest translation in response to elevated levels of amino acids in the cell. Developing similar systems that can respond to different synthetic compounds could spearhead the development of biological sensors for a variety of biotechnological applications. We have shown that the enrichment of arrest-inducing sequences is strongly correlated with their pause strength. This is important as it establishes the effectiveness of inverse toeprinting as a selection method for its subsequent use as a means to identify rare arrest sequences hidden within a complex library. While our method allows us to perform consecutive rounds of selection, it will be necessary to develop a counter-selection procedure to reduce the number of false positives after each round. In particular, we will need to address the incomplete recycling of ribosomes at stop codons that is observed currently.

The fact that inverse toeprinting is an *in vitro* method offers some advantages relative to *in vivo* approaches, in particular when it comes to identifying arrest peptides that sense small molecules. Indeed, problems may arise *in vivo* due to the inefficient uptake of small molecules into the cell, their degradation, modification or their accelerated efflux out of the cell. An additional advantage of our *in vitro* method is that it enables us to dissect molecular processes in isolation by giving us direct control over reaction conditions, as we have shown for the EF-P susceptibility of the XP(W) motifs in the presence of Ery. Being able to remove certain molecular processes from their cellular context is a double-edged sword however, and care must be taken to validate results obtained by inverse toeprinting with *in vivo* assays. For the systems studied here, the correlation between *in vitro* and *in vivo* data is generally good, but the small discrepancies observed could be due to the NNS_15_ library’s intrinsic focus on the early cycles of translation.

Finally, our method may also be used as a general tool to study sequence-dependent translational events. For example, inverse toeprinting may be modified to study the sequence-dependent impact of various protein factors on the speed of translation (e.g. signal recognition particle, SecYEG translocon, translation factors like EF-4/LepA), providing a complementary approach to ribosome profiling. The concepts presented here may further be extended to operate *in vivo* or in eukaryotes, paving the way for numerous applications.

## Methods

### Accession codes

National Center for Biotechnology Information Short Read Archive: SRP140857.

## Acknowledgments

We thank A. Malhotra for providing RNase R, N. Vazquez-Laslop and A.S. Mankin for providing the pERMZα reporter plasmid, *E. coli* TB1 cells and oleandomycin, and A.C. Seefeldt for purifying fully modified EF-P. We thank A. Buskirk for providing the pause scores for PP-containing motifs calculated from ribosome profiling data. C.A.I., B.S. and G.S. received funding for this project from the European Research Council (ERC) under the European Union’s Horizon 2020 research and innovation program (Grant Agreement No. 724040). C.A.I. is an EMBO YIP and the recipient of a Marie Curie career integration grant (PCIG14-GA-2013-631479). D.D. is funded by Inserm. B.S. and C.A.I. received funding from the Fondation pour la Recherche Médicale (AJE201133), the Région Aquitaine (2012-13-01-009) and a Chaire d’Installation from the excellence initiative (IdEx) of the University of Bordeaux awarded to C.A.I.

## Author contributions

B.S., D.D. and C.A.I. designed the experiments. B.S. and G.S. performed the experiments. B.S., G.S., D.D and C.A.I. analyzed the data. B.S., D.D. and C.A.I. drafted and revised the manuscript. All authors reviewed the manuscript and provided comments.

## Competing financial interest

The authors declare no competing financial interests.

## Online methods

### 1.1 Method overview

The DNA template encoding a T7 RNA promoter, followed by a ribosome binding site, a potential arresting peptide, a fixed ‘spacer’ region of four codons, two TGA stop codons and an *EcoRV* restriction site is generated by PCR and subsequently transcribed *in vitro* using T7 RNA polymerase with an excess of thio-phosphate-GMP, which can only be incorporated at the 5’ end of the mRNA. In the next step biotin-maleimide is coupled to the 5’ thiol group on the mRNA. The 3’ polyA-tail needed for efficient degradation by RNase R is added using Poly-A polymerase. ~5 pmol of this 5’-biotinylated and 3’-polyadenylated mRNA is then used as a template for *in vitro* translation using a PURExpress kit (NEB) from which RF-2 is omitted to prevent the release of ribosomes that translate beyond the spacer sequence. Using an NNS (a<u>**N**</u>y a<u>**N**</u>y <u>**S**</u>trong - i.e. G or C) library ensures that no other UGA stop codons should appear in the variable region. In contrast, ribosomes that reach the UAG stop codons found within the variable region can be released using RF-1.

Ribosomes engaged in translation of the mRNAs can either stall on the variable coding region if it encodes an arrest peptide or can translate until they reach the stop codon downstream of the spacer. RNase R is then used to degrade mRNAs from their 3’ end in the presence of 50 mM Mg^2^+, which inactivates and stabilizes ribosomes on the mRNA, thus ensuring that the upstream coding region is protected. Transcripts without ribosomes are degraded in this step. After this step, mRNAs are subjected to a phenol-chloroform extraction in order to ensure the inactivation and removal of RNase R and of the ribosomes. The mRNAs are purified out via their 5’-biotin using streptavidin-coupled Dynabeads. In the next step a DNA oligonucleotide linker is attached enzymatically to the 3’ end of the mRNA. This linker encodes the fixed ‘spacer’ region, followed by three TGA stop codons, one in each reading frame, followed by a restriction site. Two different linker oligonucleotides were used in this study, one encoding an *ApoI* restriction site and one encoding an *EcoRV* restriction site. Adding the linker generates a 3’ end of known sequence needed for reverse transcription of the mRNA. After second strand synthesis the double stranded cDNA is treated with the restriction enzyme encoded in the DNA template (*EcoRV* for odd rounds of selection, *ApoI* for even rounds). Ribosomes that reach the stop codon (and thus translated a sequence that does not induce stalling) protect the restriction site from RNase R degradation, thus allowing restriction enzymes to cut these DNAs and prevent their amplification in the following PCR step. Double stranded cDNAs derived from mRNAs coding for peptides that arrested the ribosome do not contain the restriction site and are consequently amplified in the ensuing PCR. The *EcoRV* and *ApoI* sites have a stretch of at least 4 A/Ts and thus cannot occur within the NNS_15_ region. PCR products from this step serve as templates for either another round of inverse toeprinting through the addition of a T7 promoter, or for NGS library generation and deep sequencing.

### 1.2 Experimental Procedures

DNA and RNA products at various points in the inverse toeprinting protocol were analyzed on 9% acrylamide (19:1) TBE (90 mM Tris, 90 mM boric acid, 2 mM EDTA) gels and stained with SyBR Gold (Invitrogen). Inverse toeprints were excised from 12% acrylamide TBE gels using a clean scalpel. Gels to analyze RNA were run under denaturing conditions (8M urea in the gel). All reactions were performed using molecular biology grade H_2_O (Millipore). Oligonucleotides used in this study are listed in **Supplementary Tables 2–5**.

#### DNA template generation

All DNA templates for inverse toeprinting were generated by PCR with Phusion DNA polymerase, using an oligonucleotide encoding the variable region in combination with oligonucleotides *T7_RBS_ATG_f* and *Stop_EcoRV_r* as templates. Amplification was performed using oligonucleotides *T7_f* and *EcoRV_r* oligonucleotides in 10-fold excess. PCR conditions included an annealing temperature of 64°C and 20 cycles of amplification. DNA templates were generated using a total of 5 pmol *ermBL_deep_mutated* oligonucleotide or 10 pmol *NNS15* oligonucleotide, respectively. PCR products were purified using a PCR purification kit (Qiagen) according to the manufacturer’s instructions and were used as templates for *in vitro* transcription.

#### *In vitro* transcription

The DNA template encodes a T7 promoter followed by an optimized Shine-Dalgarno sequence, according to the instructions of the NEB PURExpress system handbook. *In vitro* transcription was performed using T7 RNA polymerase (Promega) in a buffer containing 80 mM Tris-HCl, 24 mM MgCl_2_, 2 mM spermidine, 40 mM DTT, pH 7.6 in the presence of 7.5 mM ATP, CTP and UTP, 0.75 mM GTP (CTP, UTP GTP from Jena Bioscience) and 6.75 mM Thio-Phosphate-GMP (Genaxxon). In the first round of inverse toeprinting 10 ng/μL of DNA template were used, for further rounds the amount was reduced to 1 ng/μL. *In vitro* transcription was performed at 37°C for 2-3 hours. mRNA was purified by phenol-chloroform extraction, washed three times with chloroform and precipitated using 0.1 volumes of NH_4_-acetate (10 M) and 1 volume of isopropanol. In order to remove unincorporated nucleotides the recovered mRNA was subsequently washed through Amicon membrane centrifugal concentrators with a molecular weight cutoff (MWCO) of 30 kDa (Millipore) until the flow-through was free of unincorporated nucleotides (as determined by NanoDrop measurements). The final concentration of mRNA was determined using the NanoDrop.

#### Biotinylation

Biotinylation was performed using a 1000-fold excess of biotin-maleimide (Vectorlabs) over mRNA 5’ ends. According to the manufacturer’s instructions, the biotin-maleimide was dissolved in dimethylformamide (DMF). 600 pmol mRNA were mixed with 600 nmol biotin-maleimide in 100 mM in Bis-Tris-acetate buffer pH 6.7 and incubated at room temperature for 3 hours. Unincorporated biotin was removed by washing the mRNA three times with H_2_O (molecular biology grade, Millipore) in an Amicon membrane centrifugal concentrator with a MWCO of 30 kDa (Millipore). mRNA was recovered and biotinylation efficiency was analyzed using a Dot Blot.

#### Dot Blot

H+ bond membrane (GE healthcare) was treated with 6x SSC buffer (900 mM NaCl, 90 mM Na_3_-citrate, pH 7.0) for 10 minutes and dried briefly between two pieces of Whatman paper. Samples and standard were diluted in 6x SSC buffer to 0.5, 1.0, 2.5 and 5.0 μM and 1 μL of each dilution was pipetted onto the prepared membrane. The membrane was then baked for two hours at 80°C to attach the mRNA to the membrane. The membrane was subsequently blocked in 2.5 % dry milk solution in TBS-T (50 mM Tris-HCl, 150 mM NaCl, 0.05 % (v/v) Tween-20, pH 7.5) for 1 h at room temperature. The milk solution was removed and the membrane was incubated with Streptavidin-alkaline phosphatase antibody (Promega) in a 1:1000 dilution in TBS-T for 1 h at room temperature. Unbound antibody was removed by washing three times with TBS-T buffer. Detection was performed using the NBT/BCIP detection kit (Promega) in alkaline phosphatase buffer according to the manufacturer’s instructions. The detection reaction was stopped by two washes with TBS-T buffer and a picture of the membrane was taken immediately on BioRad Imager. The biotinylation efficiency was estimated from the intensity of the sample dots compared to the intensity of the standard dots.

#### Poly-adenylation of the mRNA 3’ end

Poly-adenylation of the biotinylated mRNA was performed using Poly-A-polymerase (NEB) using the supplemented buffer. The ratio of mRNA 3’ ends to ATP molecules was chosen to be 1:100. The reaction was incubated at 37°C for 2-3 hours and poly-adenylation efficiency was assessed by denaturing PAGE (9%). Polyadenylated mRNA was purified using phenol-chloroform extraction, washed three times with chloroform and precipitated with NH_4_-acetate-isopropanol with 0.5 μL GlycoBlue (Thermo Fisher). Unincorporated ATP was removed by washes using Amicon membrane centrifugal concentrators with a MWCO of 30 kDa (Millipore).

#### RNase R activity

Purified RNase R^29^ was provided by Dr. Arun Malhotra (University of Miami) and was used at 1 mg/mL stock solution. 5 pmol of mRNA were used to test the degradation efficiency of 2 μL RNase R on every batch of mRNA in a buffer containing 50 mM HEPES-KOH, 100 mM K-glutamate, 50 mM Mg-acetate, 1 mM DTT, pH 7.5. Time points were taken directly into RNA loading dye (95% formamide, 250 μM EDTA, 0.25% (w/v) bromophenol blue, 0.25 % (w/v) xylene cyanol) before addition of the enzyme and after 5, 10 and 30 minutes of incubation at 37°C. The samples were analyzed by denaturing PAGE (9%), stained with SyBR Gold to monitor mRNA degradation.

#### Inverse toeprinting

PURExpress Δ RF123 kit (NEB) was used to perform *in vitro* translation. ~5 pmol of 5’-biotinylated and 3’-polyadenylated mRNA were used as a template. Antibiotics (Ery, Ole) were supplemented at a final concentration of 50 μM in 10 μL reactions. RF-1 and RF-3 were added to the translation reaction according to the manufacturer’s instructions. Translation was performed at 37°C for 30 minutes before the samples were placed on ice and 10 μL ice-cold Mg^2^+ buffer (50 mM HEPES-KOH, 100 mM K-glutamate, 87 mM Mg-acetate, 1 mM DTT, pH 7.5) was added to the reactions, thus increasing the Mg^2+^ concentration to 50 mM. 2 μL of RNase R (1 mg/mL) were added, followed by an additional incubation for 30 minutes at 37°C for RNase R-mediated mRNA degradation. Ribosome-protected mRNA was purified by phenol-chloroform extraction, washed three times with chloroform and precipitated using NH_4_-acetate-isopropanol. RNA was recovered by centrifugation at full speed for 30 minutes at 4°C and resuspended in 50 μL 1x BWT buffer (5 mM Tris-HCl, 0.5 mM EDTA, 1 M NaCl, 0.05 % (v/v) Tween-20, pH 7.5).

#### mRNA purification with Dynabeads

5 μL M-280 streptavidin Dynabeads (Life Technologies) were prepared for each sample by washing three times with 1x BWT buffer in DNA loBind tubes (Eppendorf) and resuspended in 50 μL of the same buffer. Dynabeads and purified RNA from the previous step were combined in these tubes and incubated on a tube rotator for 15 minutes at room temperature to allow binding of the biotinylated mRNA to the streptavidin beads. After incubation, the beads were collected using a magnet and the supernatant was discarded. The beads were washed two times with 1x BWT buffer to remove unincorporated RNA, followed by two washes with H_2_O to remove the 1x BWT buffer. Beads were resuspended in 4.5 μL H_2_O.

#### Linker ligation

The beads from the previous step were combined with 10 pmol (1 μL) of the desired linker (*3’_linker_ApoI* or *3’_linker_EcoRV* depending of the round) plus 3 μL PEG 8,000, 1 μL PNK buffer and 0.5 μL T4 RNA ligase 2, truncated (all NEB). Incubation was performed on a tube rotator for 2 hours at room temperature. After incubation the beads were washed three times with H_2_O to remove unincorporated linker oligonucleotide and were resuspended in 12 μL H_2_O.

#### Reverse transcription

12 μL beads were combined with 1 μL *Linker_r* oligonucleotide (2 μM) and 1 μL dNTPs (NEB, 10 mM per dNTP) and incubated at 65°C for 5 minutes to allow annealing of the *Linker_r* oligonucleotide to the linker. After annealing 4 μL of 1^st^ strand buffer, 1 μL of 100 mM DTT and 1 μL of superscript III (all Invitrogen) were added and the reaction incubated at 55°C for 30 minutes to allow reverse transcription of the Dynabead-bound mRNA.

#### PCR on cDNA, restriction digestion

Reverse transcribed cDNA was used without further purification as template for PCR. In order to generate double stranded DNA for restriction digestion, a fill-up reaction was performed using *cDNA_f* oligonucleotide and the reverse transcribed cDNA (10 s denaturation, 5 s annealing at 42°C and 30 s elongation at 72°C). The resulting dsDNA was combined with 1 μL of the respective restriction enzyme and the sample incubated at 37°C for 1 hour. To amplify undigested DNA *Linker_r* oligonucleotide was added and a PCR performed with 8-14 cycles (Denaturation at 98°C for 10 s, annealing at 42°C for 5 s, elongation at 72°C for 10 s). The number of PCR cycles giving the best results was used for further purification of the cDNA.

#### Purification of DNA fragments of interest after PCR

Wild-type and *ermBL* samples were purified using a homemade electro-elution device. PCR products were analyzed by TBE-PAGE (12%) and bands of interest were excised from the gel using a clean scalpel. Gel pieces were crushed through a 5 mL syringe into 50 mL Falcon tubes whose base (1-2 mL) had been cut off and covered with Parafilm. The crushed gel pieces were then embedded into a new 9% acrylamide TBE gel inside the cut Falcon tube (approx. 8 mL of gel solution). After polymerization, DNA was eluted from the gel by filling the Falcon tube with TBE buffer (upper buffer reservoir) and hanging the Falcon tube into a beaker filled with TBE buffer (lower buffer reservoir, ice-cooled). DNA was eluted into the lower buffer reservoir by placing clean electrodes into the two buffer reservoirs elution and by applying a current of 10W per gel (2 gels max) for 30 minutes. The buffer from the lower reservoir was recovered and the eluted DNA was concentrated using Amicon membrane centrifugal concentrators with a MWCO of 30 kDa (Millipore). DNA was precipitated by addition of NH_4_-acetate isopropanol and 0.5 μL GlycoBlue (Thermo Fisher) and incubation at −20°C for 1 hour.

The NNS_15_ library samples were purified using a modified method from Ingolia *et al*.^30^. After cutting out the bands of interest the gel pieces were crushed through a 5 mL syringe into 15 mL Falcon tubes and 7.5 mL of gel elution buffer (10 mM Tris-HCl, pH 8.0, 500 mM Na-acetate, 0.5 mM Na-EDTA) were added. The tubes were incubated on a tube rotator at room temperature overnight. Gel debris was separated from the buffer by filtering through 0.22 μm centrifugal filters (Millipore). Each sample was then concentrated to ~250 μL using a SpeedVac. The eluted DNA was precipitated using 0.1 volume NH_4_-acetate and 2.5 volumes ethanol with 0.75 μL GlycoBlue (Thermo Fisher) and incubation at 242:80920°C for 1 hour.

After precipitation, DNA was recovered by centrifugation in a tabletop centrifuge at full speed for 30 minutes at 4°C. The supernatant was carefully removed by pipetting and the DNA pellet briefly dried in the SpeedVac for 10-15 minutes. The cDNA was then resuspended in 15 μL H_2_O (molecular biology grade) and used as a PCR template for the addition of the T7 promoter sequence or the NGS adapters.

#### Addition of T7 promoter for another round of inverse toeprinting

The purified cDNA from the previous step was used as a template for PCR in combination with *T7_RBS_ATG_f* (1 μM), encoding the T7 promoter needed for *in vitro* transcription, *T7_f* and *Linker_r* oligonucleotides (10 μM each). 8-14 cycles of PCR (64°C annealing temperature) were used and amplified DNA was purified using Qiagen PCR purification kit. The concentration of purified DNA was determined using a NanoDrop.

#### Addition of NGS adaptor to purified cDNA

Long NGS adaptor oligonucleotides contain the Illumina TruSeq adapter sequences followed by 18 nt complementary to the 5’ or 3’ region of the cDNA. The reverse NGS oligonucleotides also encode barcode sequences for multiplexing according to the TruSeq v1/v2/LT protocol (Illumina). The adaptors were added to the cDNA by PCR (8-14 cycles) using the long oligonucleotides (20-26) in 1 μM stock solutions and the short amplifying oligos (18, 19) in 100 μM stock concentration. PCR products were purified using Qiagen PCR purification kit. The size and concentration of the fragments obtained were analyzed using a 2100 Agilent Bioanalyzer with the DNA 1000 kit.

#### Preparation of mRNA template library for NGS

In order to prepare the mRNA library for NGS 5 pmol of mRNA were reverse transcribed using Superscript III (Invitrogen) using an NGS adapter (27/28) containing a stretch of 18 nt reverse complementary to the fixed ‘spacer’ region in the 3’ end of the mRNA. The resulting cDNA served as template for subsequent PCR using the forward (20) and reverse adapter (27/28) in 1 μM stock concentration and the short amplifying oligonucleotides in 100 μM stock concentration. PCR products were purified using a Qiagen PCR purification kit. The size and concentration of the fragments obtained were analyzed using a 2100 Agilent Bioanalyzer with the DNA 1000 kit.

#### Next generation sequencing

Next generation sequencing was performed by the Tufts Genomics Core Facility (TUCF) in Boston, USA on an Illumina HiSeq 2500 system in rapid run mode with 150 PE read.

#### Toeprinting

Toeprinting to test novel arrest sequence motifs was performed using the PURExpress ΔRF1,2,3 kit (NEB). DNA templates were generated by PCR using *T7_RBS_ATG_f, TP_3’_spacer_r* and *TP_NV1_r* oligonucleotides (all at 1 μM) in combination with the oligonucleotide encoding the sequence to be tested. These oligonucleotides served as templates and were amplified using the *T7_f* and *TP_NV1_r_short* oligonucleotides (100 μM) with Phusion DNA polymerase. PCR products were purified using the Qiagen PCR purification kit and eluted with H_2_O (molecular biology grade). Ery or Ole were dried into the tube to yield a final concentration of 50 μM in the 5 μL toeprinting reaction. 1 pmol of DNA template was combined with 2 μL of solution A and 1.5 μL solution B of the PURExpress system. The reaction was incubated at 37°C for 15 minutes prior to addition of 1 μL of the 5’-Yakima Yellow labeled NV1 probe^31^ (2 μM) and the reaction was incubated for another 5 minutes at 37°C. Reverse transcription was performed with 0.1 μL dNTPs (10 mM each dNTP), 0.4 μL PURE system buffer and 0.5 μL AMV RT (Promega) and 20 minutes incubation at 37°C. After generation of the Yakima Yellow-labeled cDNA the mRNA was degraded by addition of 0.5 μL 10 M NaOH and incubation at 37°C for 20 minutes. The samples were neutralized with 0.7 μL 7.5 M HCl. 20 μL toeprint resuspension buffer (300 mM Na-acetate pH 5.5, 5 mM EDTA, 0.5% SDS) and 200 μL PNI buffer were added to each sample and cDNA was purified using the Qiagen nucleotide removal kit. The cDNA was eluted using 50 μL of H_2_O (molecular biology grade). cDNA was dried into the tube using a SpeedVac and resuspended in 6 μL toeprint loading dye (95% formamide, 250 μM EDTA, 0.25 % bromophenol blue). Samples were denatured at 95°C for 5 minutes prior to loading onto a 7.5 % polyacrylamide TBE sequencing gel containing 8 M urea. The gel was run at 40 W and 2000 V for 2.5 hours. Yakima-Yellow labeled cDNAs were detected using a Typhoon Gel Scanner in fluorescent mode.

#### Disc diffusion assay

We used the LacZα-based *in vivo* system described by Bailey and coworkers^26^. Using oligonucleotides 39-50 we generated by PCR several plasmids in which we replaced the first 10 codons of the encoded *ermCL* with the sequence of interest, thus maintaining the regulatory region of *ermCL* and *ermC*. We transformed these plasmids into chemically competent *E. coli* TB1 cells. Transformants were grown for 6 hours in LB media with 1 mM isopropyl beta-D-1-thiogalactopyranoside (IPTG) at 37 °C to an OD_600_ of 1.5. For the MAXPW motifs, 50 μL of IPTG (100 mM) were added to LB-agar plates containing ampicillin (100 μg/mL) and streptomycin (50 μg/mL) prior to addition of 100 μL cells. Whatman filter discs soaked with 10 μL 5-bromo-4-chloro-3-indolyl-β-D-galactopyranoside (X-gal) (50 μM) and either water or 50 μg Ery were placed onto the agar plates. For the ErmBL constructs, 50 μL of IPTG (100 mM) and 50 μL of X-gal (50 μM) were added to LB-agar plates containing ampicillin (100 μg/mL) and streptomycin (50 μg/mL) prior to addition of 100 μL cells. Whatman filter discs soaked with water or 50 μg Ery or Ole were placed onto the agar plates. Agar plates were incubated at 37 °C for 24h and pictures were taken immediately after the incubation period.

#### Expression and purification of Elongation Factor P

Expression of EF-P was performed together with the expression of the modification enzymes as described previously^32^ in *E. coli* BL21 Gold and expression was induced using 1 mM IPTG at an OD_600_ of 0.6. Harvested cells were resuspended in lysis buffer (50 mM HEPES KOH, pH 7.6, 10 mM MgCl_2_, 1 M NH_4_Cl) and sonicated. Cell debris was removed by centrifugation (45 minutes, 40,000 x g, 4 °C) and the clarified lysate was mixed with cobalt-agarose (Sigma) and incubated on a tube rotator at 4 °C for 1 hour. The cobalt-agarose was washed with lysis buffer and the protein was eluted with lysis buffer containing 250 mM imidazole. The eluate was concentrated using centrifugal concentrators with a MWCO of 10 kDa (Millipore). In order to exchange the buffer to protein storage buffer (20 mM HEPES KOH, pH 7.6, 10 mM MgCl_2_, 50 mM KCl, 50 mM NH_4_Cl) gel filtration was performed on a Superdex 75 using an NGC medium pressure liquid chromatography system (BioRad). Eluate fractions were analyzed by SDS-PAGE and protein-containing fractions were pooled and concentrated to a final concentration of 20 mg/mL using centrifugal concentrators with a MWCO of 10 kDa (Millipore). The activity of the purified EF-P was assessed by toeprinting using a DNA template encoding the sequence MMHHHHHHRPPPI. Addition of EF-P to a final concentration of 10 μM efficiently rescued ribosomes stalled on the poly-proline motif.

### 1.3 Data analysis

Unless it is indicated otherwise, data analysis was carried out using a series of custom scripts written in-house in Python, which relied upon the use of the Biopython package^33^.

#### Read assembly and trimming

Read pairs were assembled using PEAR v0.9.10^34^ on a Mac Book Pro with a 2.7 GHz Intel Core i7 processor and 16 GB 1600 MHz DDR3 memory, with the maximal proportion of uncalled bases in a read set to 0 (–u option) and the upper bound for the resulting quality score set to 126 (–c option).

Regions immediately upstream of the start codon and downstream of the point of cleavage by RNase R were removed using a modified version of the *adaptor_trim.py* script written by Brad Chapman (https://github.com/chapmanb/bcbb/blob/master/align/adaptor_trim.py). The 5’ flanking region was defined as GTATAAGGAGGAAAAAAT, while the 3’ flanking region was GCGATCTCGGTGTGATG for the NNS_15_ and ErmBL libraries, and GGTATCTCGGTGTGACTG for all other samples. A maximum of 2 mismatches within each of these flanking regions was tolerated, while all other reads were discarded. Trimming of the retained reads resulted in sequences with a start codon directly at the 5’ end and, in the case of samples resulting from inverse toeprinting, the site of RNase R cleavage at the 3’ end.

#### Quality filtering and selection of the region of interest

Trimmed sequences were further processed with our *process-reads.py* script. Reads featuring ≥18 ‘A’ bases within the first 22 nucleotides from the start codon were eliminated, as were reads from which the expected ‘ATG’ start codon was absent. Sequences within the region of interest were retained for further analysis, provided that each base call within this region had a Q score of 30 or more. For the NNS_15_ library sample, the region of interest spanned nucleotides 1 to 48, where nucleotide 1 is the first nucleotide of the start codon. For all samples selected after translation of the NNS_15_ library, the region of interest is defined in Fig. 1 and Supplementary Fig. 6 and covered nucleotides 24 to 47. For all ErmBL-related samples, the region of interest covered nucleotides 30 to 32 from the start codon. A summary of NGS read processing is given in **Supplementary Table 6**.

#### Analysis of RNase R cleavage in known arrest sequences

Assembled and trimmed reads for all of the known arrest sequences shown in **Fig. 1b,c** were processed using the *process-reads.py* script, with the region of interest covering nucleotides 24 to 77 and the minimum Q score for base calls within this region set to 60. We then obtained the size distribution of reads that were exact matches to the 5’ end of each of the sequences in **Supplementary Table 7**. This task was automated with our *find-exact-match.py* script.

#### Translation into amino acid sequences

Processed reads were translated using the *translate-reads.py* script. For samples that had undergone inverse toeprinting, the ribosome-protected region downstream of the A-site codon was removed prior to translation. Unique peptidic sequences were identified and the frequency of occurrence of these sequences within each sample was calculated.

#### Calculation of 3-aa motif frequencies

All possible 3-aa motifs centered on the P-site of the stalled ribosomes were identified and counted within the translated sequences, using the *process-kmers.py* script with a word size of 3. In each case, the frequency of occurrence (F_3aa_) of the 3-aa motif was

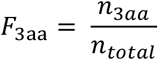

where *n*_3*aa*_ is the number of occurrences of this 3-aa motif at the C-terminus of the translated sequences and *n_total_* is the total number of processed reads in the sample.

#### Calculation of fold changes and propagation of inter-replicate errors

The fold change in 3-aa motif frequency between two samples was calculated using the *read-analyzer.py* script and was defined as *F*_Fg_/*F*_Bg_, where *F*_Fg_ is the frequency of occurrence of a sequence or 3-aa motif in the “foreground” sample and *F*_Bg_ is its frequency in the “background” sample. For the comparison between Ery-treated and untreated samples, the mean frequency of occurrence of each 3-aa motif in the presence or absence or drug was

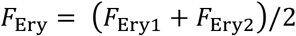

and

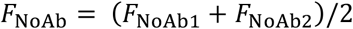

respectively.

Similarly, the errors between replicates were

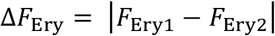

and

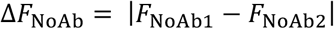

The fold change in 3-aa motif frequency upon addition of Ery was given as *F*_Ery_/*F*_NoAb_ and the combined error of the fold change in 3-aa frequency was:

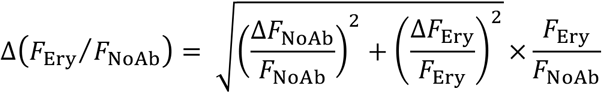

The histogram in **Fig. 3a** was built using the Δ(*F*_Ery_/*F*_NoAb_) values for all well-measured 3-aa motifs.

#### Calculation of pause strengths

Pause strengths for all 3-aa motifs were calculated using the *calculate_all_pause_strengths.py* script, according to the formula:

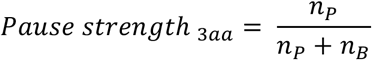

where *n_P_* is the number of reads where a ribosome is stalled on the 3-aa motif of interest and *n_B_* is the number of reads where the ribosome has translated through this motif.

---

**Supplementary Fig. 1.**
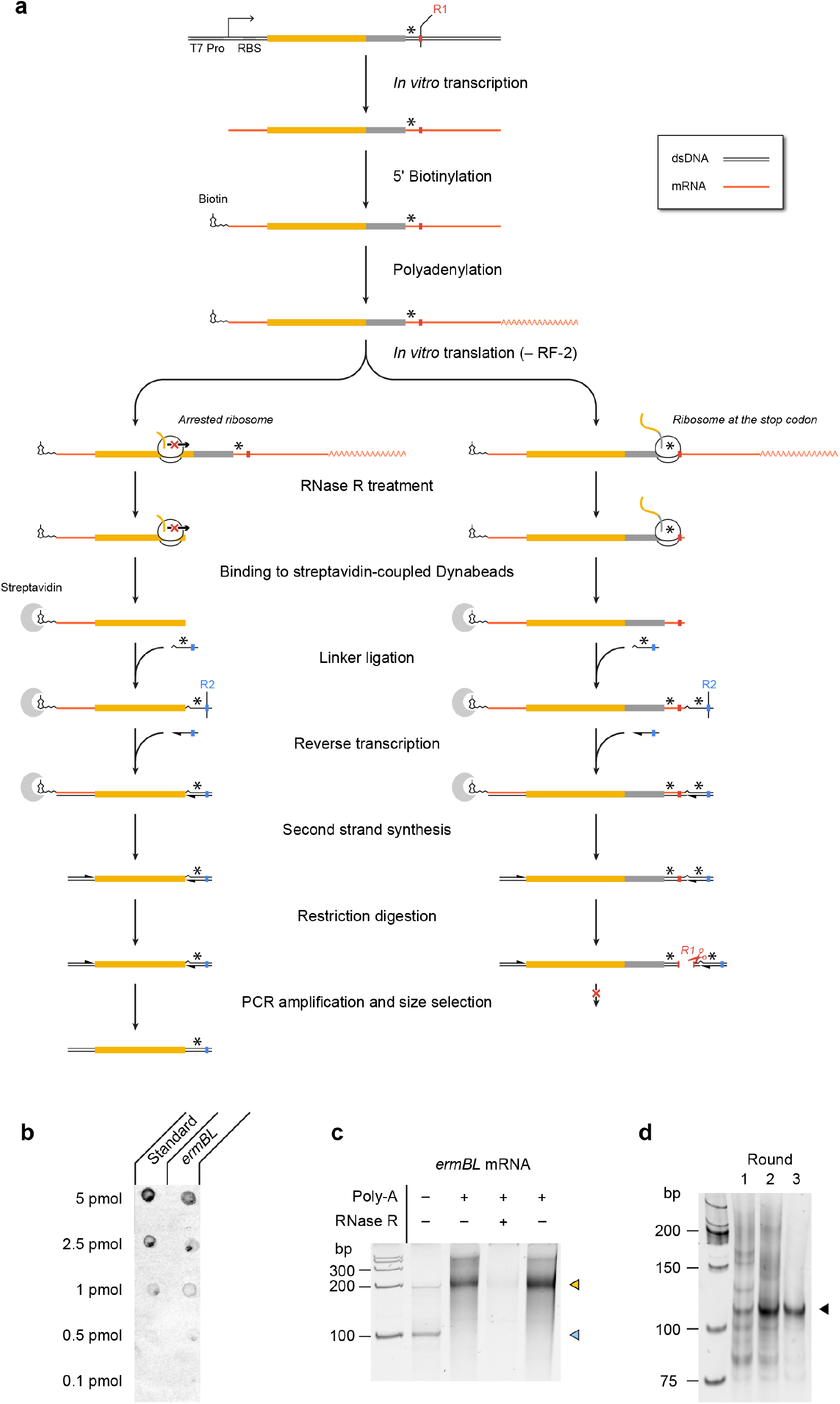
Overview of the inverse toeprinting workflow. (a) Step-by-step description of the inverse toeprinting methodology. Restriction enzymes used in odd (*EcoRV*)and even (*ApoI*) cycles are shown in red and blue, respectively. Stop codons are indicated as asterisks. (b) Efficiency of *ermBL* mRNA biotinylation measured using a dot blot assay. The standard used is a synthetic biotinylated oligonucleotide, which corresponds to 100% biotinylation efficiency. (c) Efficiency of polyadenylation and digestion by RNase R. The light blue arrow indicates untreated *ermBL* mRNA and the yellow arrow indicates polyadenylated *ermBL* mRNA. (d) Amplification of *ermBL* toeprints following 1, 2 or 3 rounds of inverse toeprinting in the presence of Ery.

**Supplementary Fig. 2.**
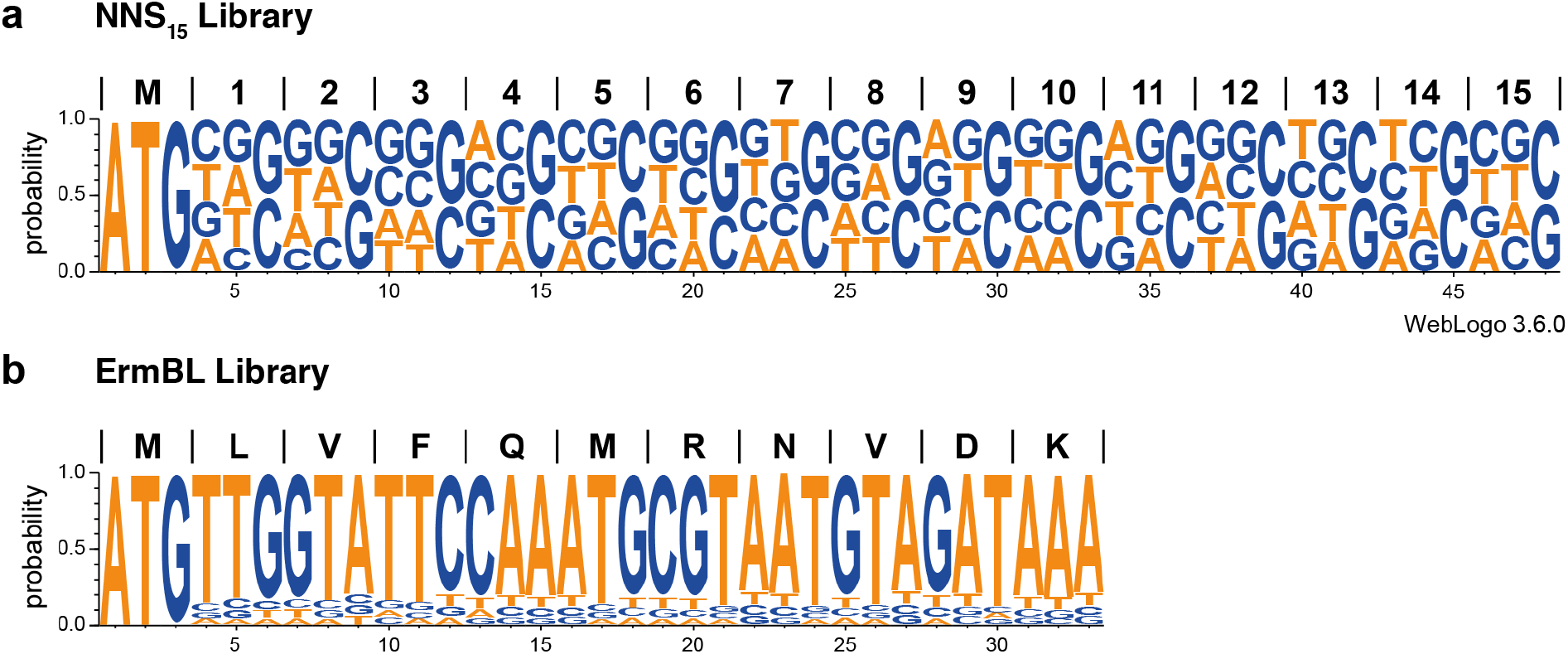
DNA template libraries used for inverse toeprinting. (a, b) WebLogos^35, 36^ obtained from 100,000 randomly chosen sequenced reads for the (a) NNS_15_ and (b) ErmBL libraries.

**Supplementary Fig. 3.**
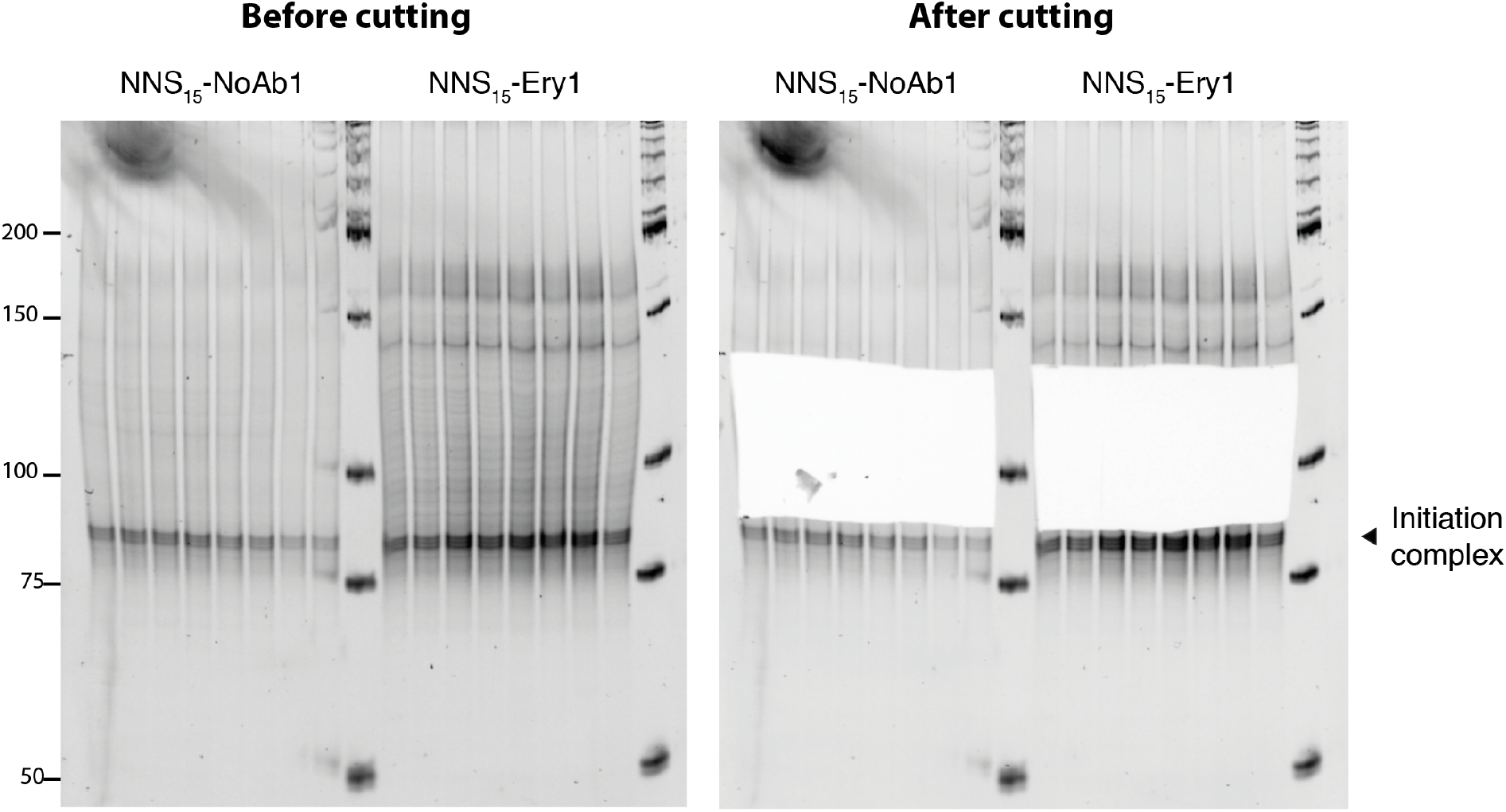
Excision of inverse toeprints from a polyacrylamide gel. cDNA obtained from inverse toeprints was amplified by PCR after second strand synthesis and *EcoRV* treatment and loaded onto a 12 % acrylamide TBE gel stained with SyBR Gold. The band indicated by a black triangle corresponds to inverse toeprints of ribosomes stalled at the initiation codon. Amplified double stranded cDNA corresponding to ~90-135 bp was excised from the gel with a clean scalpel to retain inverse toeprints where ribosomes had stalled after translating 2-15 codons. cDNA was eluted from the gel by diffusion as described in the methods section.

**Supplementary Fig. 4.**
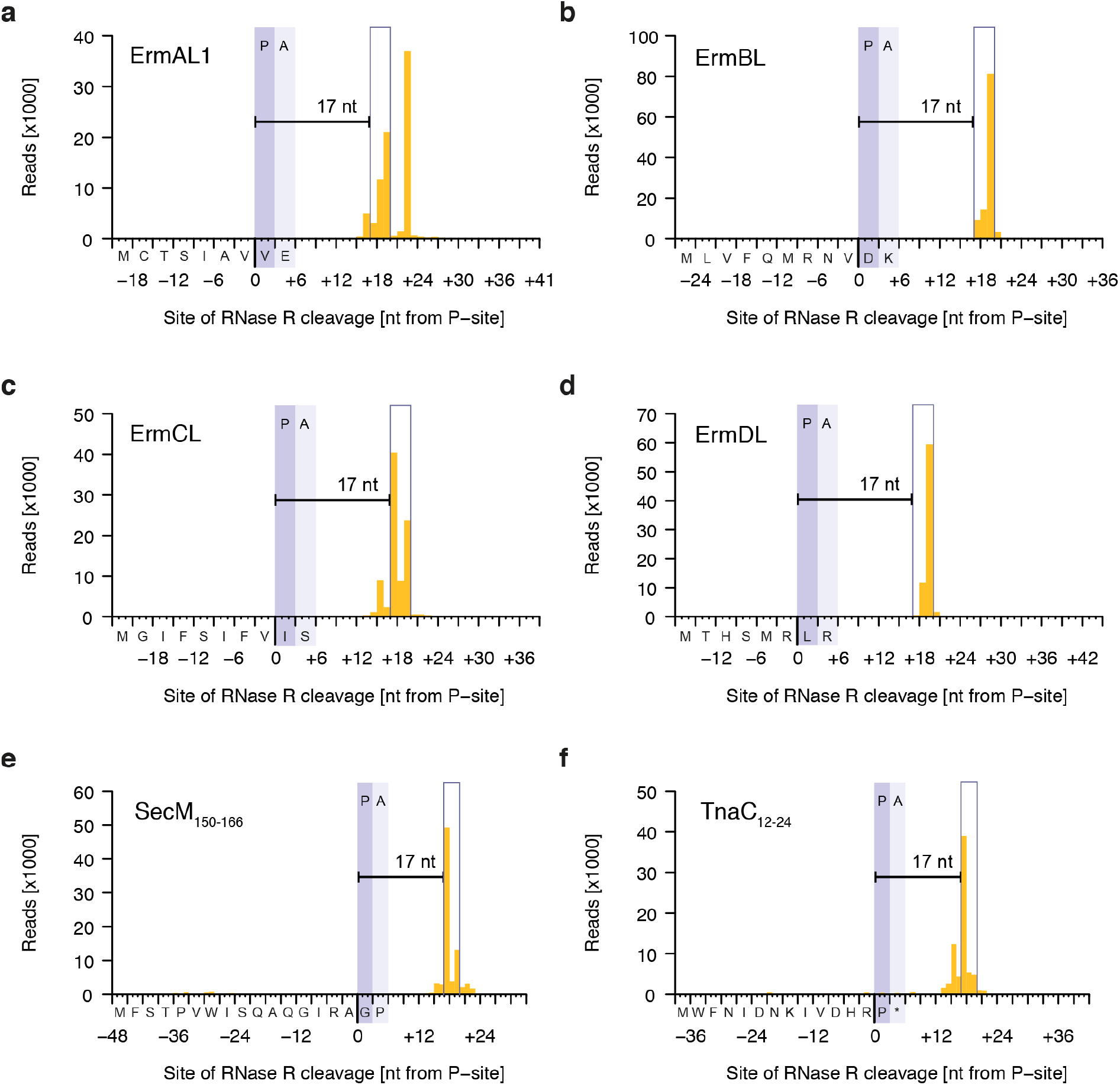
Inverse toeprints obtained for various known arrest sequences. The distance between the documented point of arrest and the site of cleavage by RNase R is shown for the known arrest peptides ErmAL1 (a), ErmBL (b), ErmCL (c), ErmDL (d), SecM_150-166_ (e) and TnaC_12-24_UAA_25_ (f). Note that ErmAL1 appears to feature a previously unidentified second arrest site.

**Supplementary Fig. 5.**
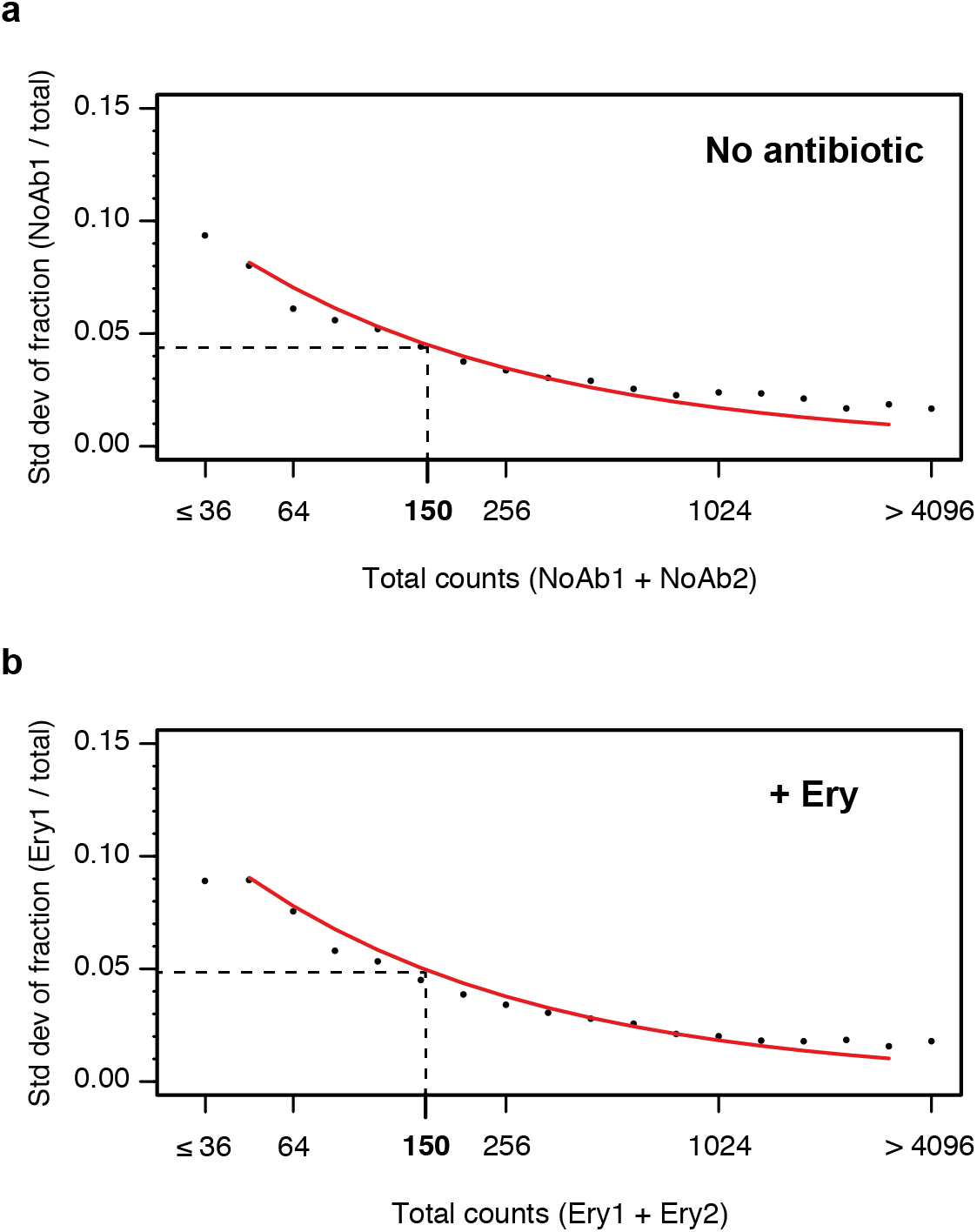
Impact of counting statistics on the quantification of errors between replicates. The reproducibility of inverse toeprinting was assessed using fully independent biological replicates. For each 3-aa motif, the fraction of the total number of reads between replicates that originated from replicates (a) NoAb1 and (b) Ery1 was calculated and binned according to the total number of reads. The standard deviation of these fractions was calculated and plotted for each bin, as indicated by black points. The red curve indicates predicted standard deviations obtained using the same number of total reads for each bin, which had been randomly partitioned into two replicates for each sample, according to probabilities that were proportional to the total number of reads in each replicate. The strong correlation between the predicted and measured standard deviations indicates that counting statistics are the main source of error. A threshold of 150 reads was chosen as the point where the standard deviation drops below 0.05.

**Supplementary Fig. 6.**
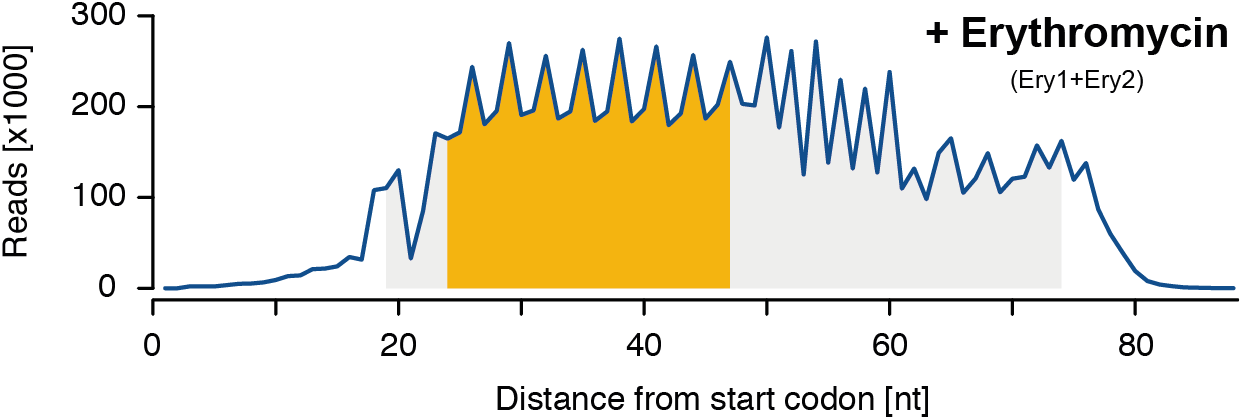
Size distribution of inverse toeprints obtained in the presence of antibiotics. Size distribution of inverse toeprints from two biological replicates with a minimum Q-score of 30 obtained from an NNS_15_ library translated in the presence of Ery (Ery1+Ery2). The fragment size ranges shaded in gray corresponds to the bands that were cut from a 12% TBE-acrylamide gel, while the ranges in yellow indicate fragments used in our analysis.

**Supplementary Fig. 7.**
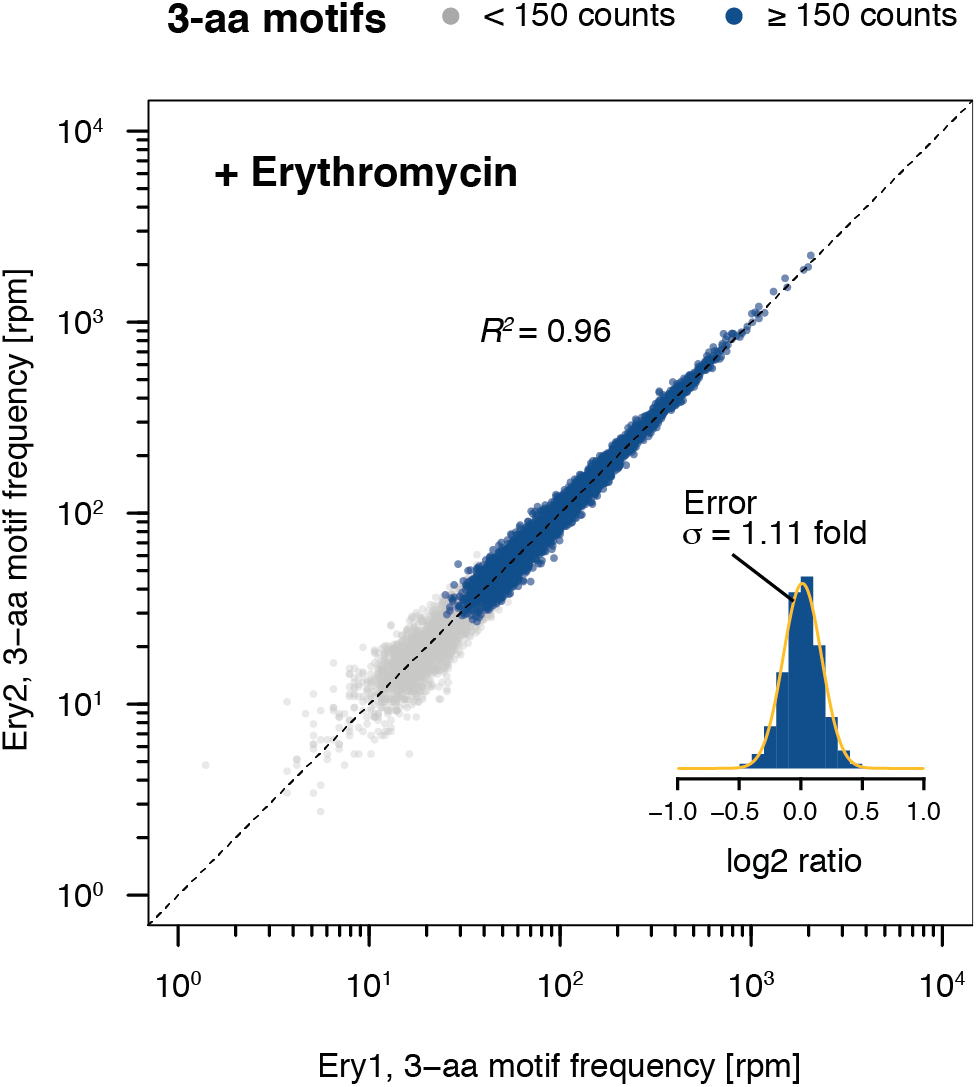
Comparison of biological replicates for Ery-treated samples. 3-aa motif frequencies in two biological replicates obtained in the presence of Ery (Ery1 and Ery2). The inset represents a histogram of log2 ratios between replicates for 3-aa motifs having low statistical counting error (i.e. with >150 counts (blue), Supplementary Fig. 5) with an overlaid normal error curve (mean = 0.01, standard deviation = 0.16 log2 units, equivalent to σ = 1.11 fold).

**Supplementary Fig. 8.**
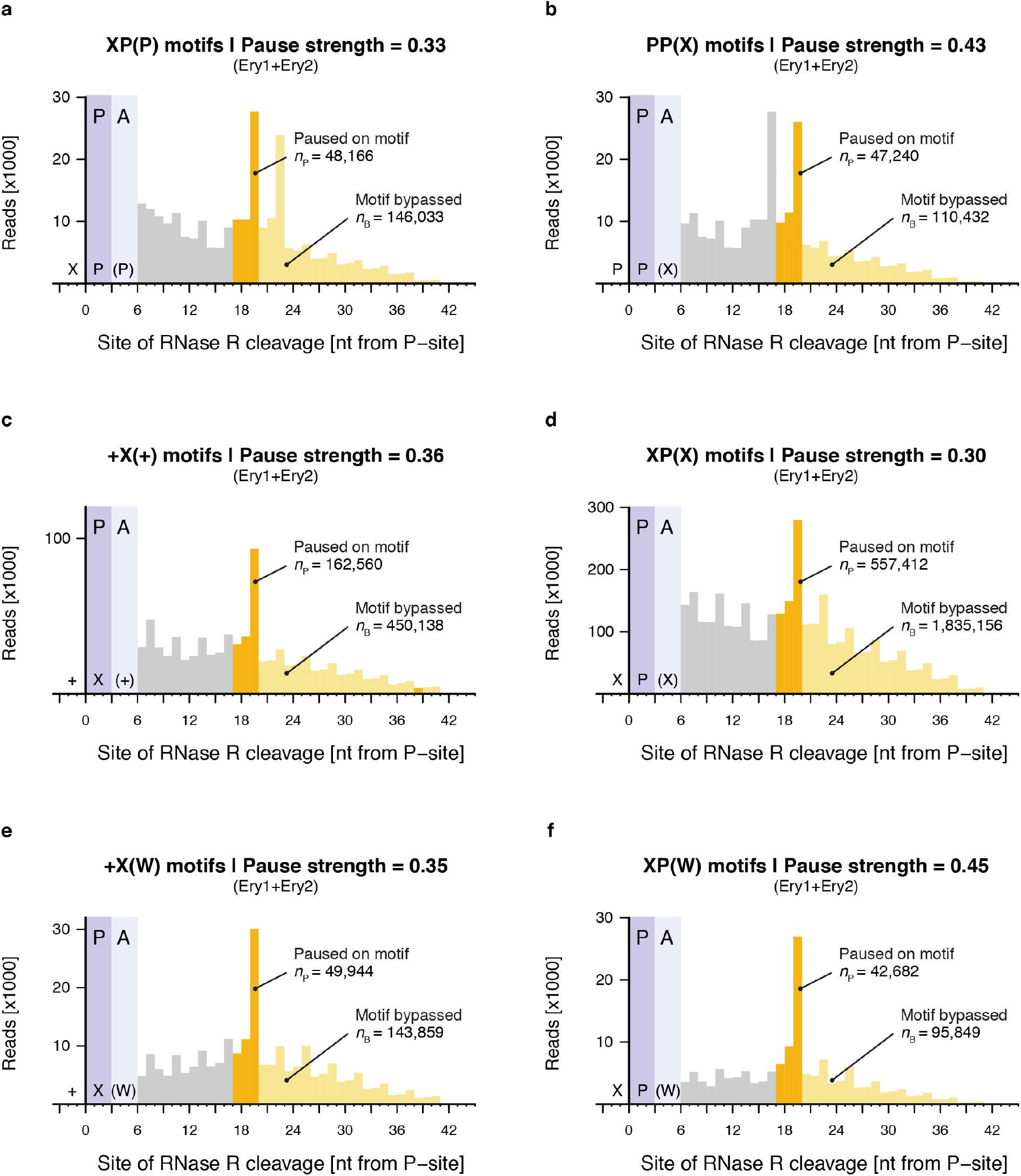
Calculation of pause strengths for various Ery-dependent arrest motifs. Histograms of the distances between (a) XP(P), (b) PP(X), (c) +X(+), (d) XP(X), (e) +X(W) and (f) XP(W) motifs and the 3’ end of inverse toeprints obtained in the presence of Ery. Both Ery replicates (Ery1 and Ery2) were combined for this analysis.

**Supplementary Fig. 9.**
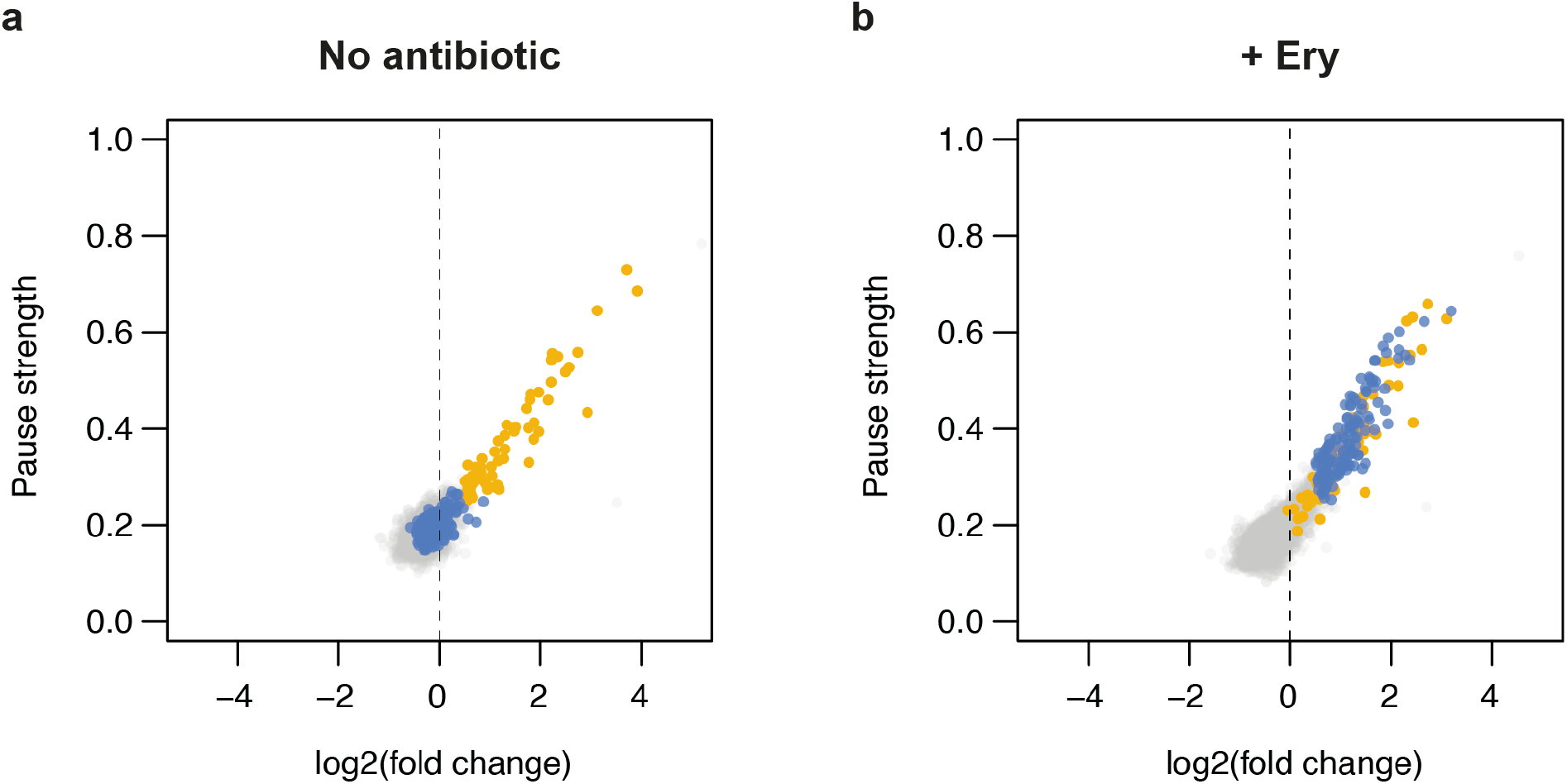
Enrichment of 3-aa motifs upon addition of Ery. (a, b) Plots of pause strength against log2(fold change) for all possible 3-aa motifs, translated in the (a) absence or (b) presence of Ery. Yellow points correspond to 3-aa motifs that induce translational pausing in the absence of antibiotic. Blue points correspond to motifs with pause strengths ≥ 0.25 in the presence of Ery and ≥ 1.5-fold greater than in the absence of drug, which show a log2(fold change) in frequency of ≥ 0.5. All other motifs are shown as gray dots.

**Supplementary Fig. 10.**
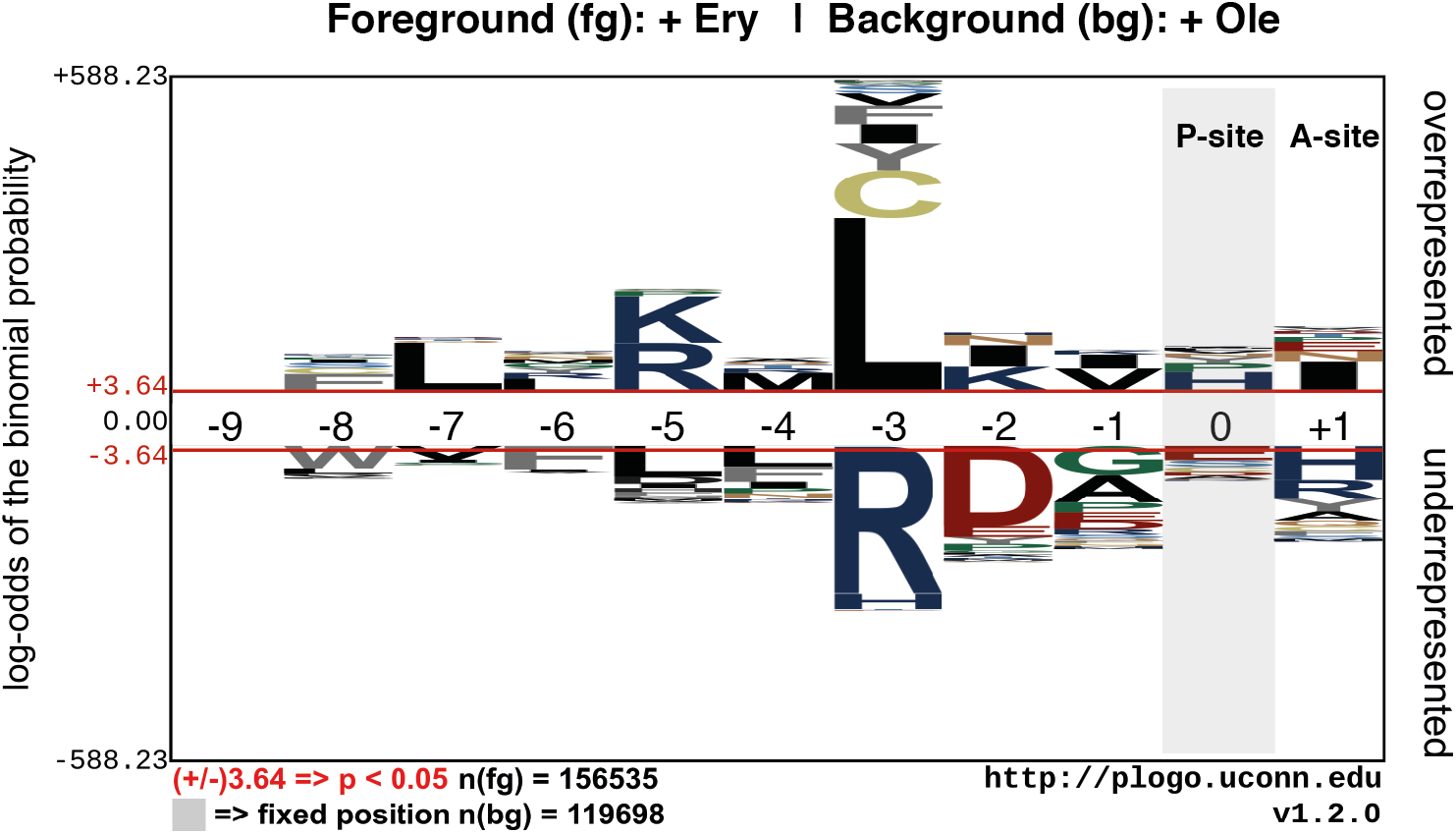
Overrepresentation or underrepresentation of amino acids at different positions in ErmBL. Unique ErmBL variants were identified from inverse toeprints obtained after translation in the presence of Ery or Ole. A comparison between these two sets of inverse toeprints was performed using the pLogo server^37^. Peptide numbering is such that residue 0 is in the P-site and +1 is in the A-site.

**Supplementary Table 1.**
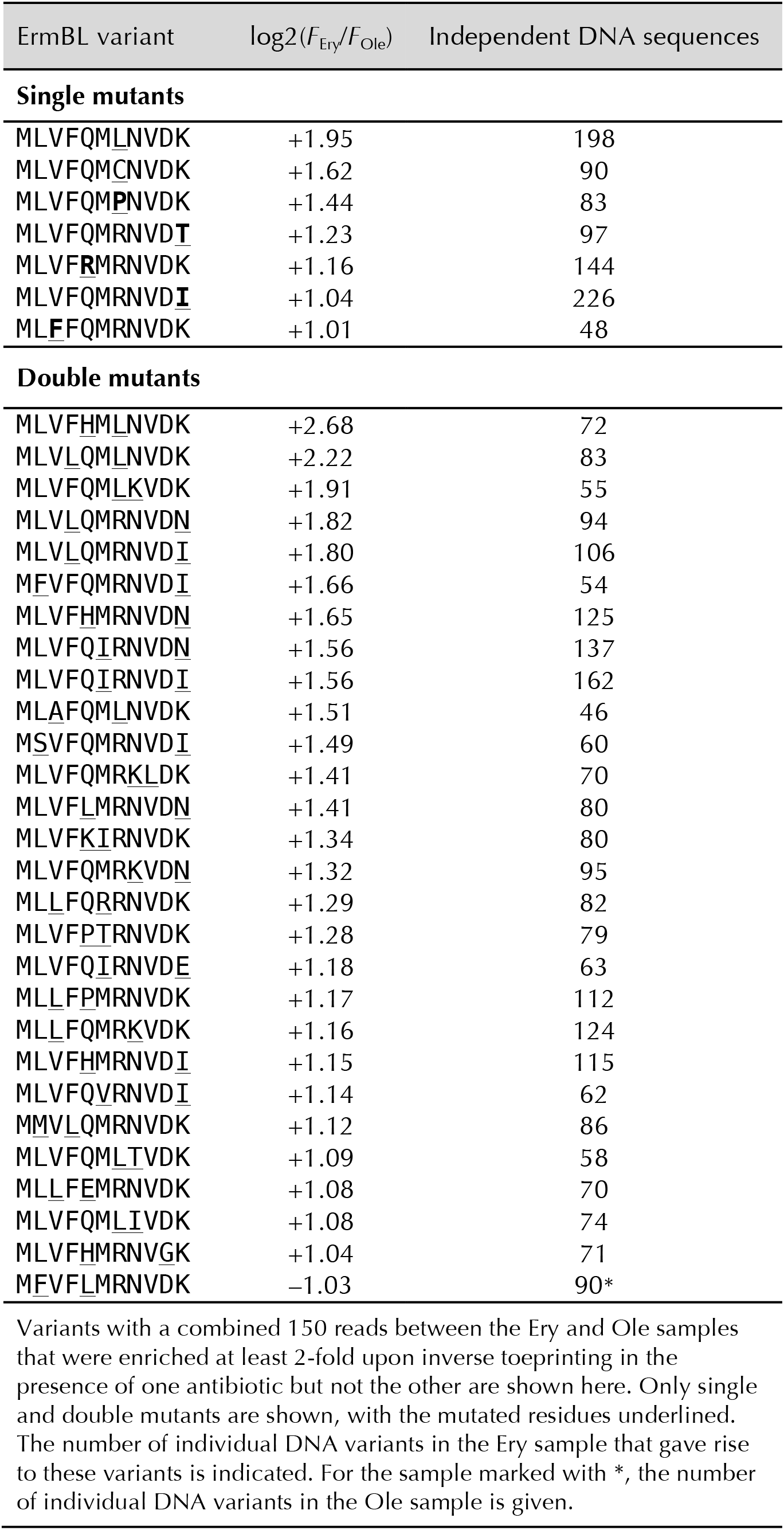
ErmBL variants that discriminate between Ery and Ole

**Supplementary Table 2.**
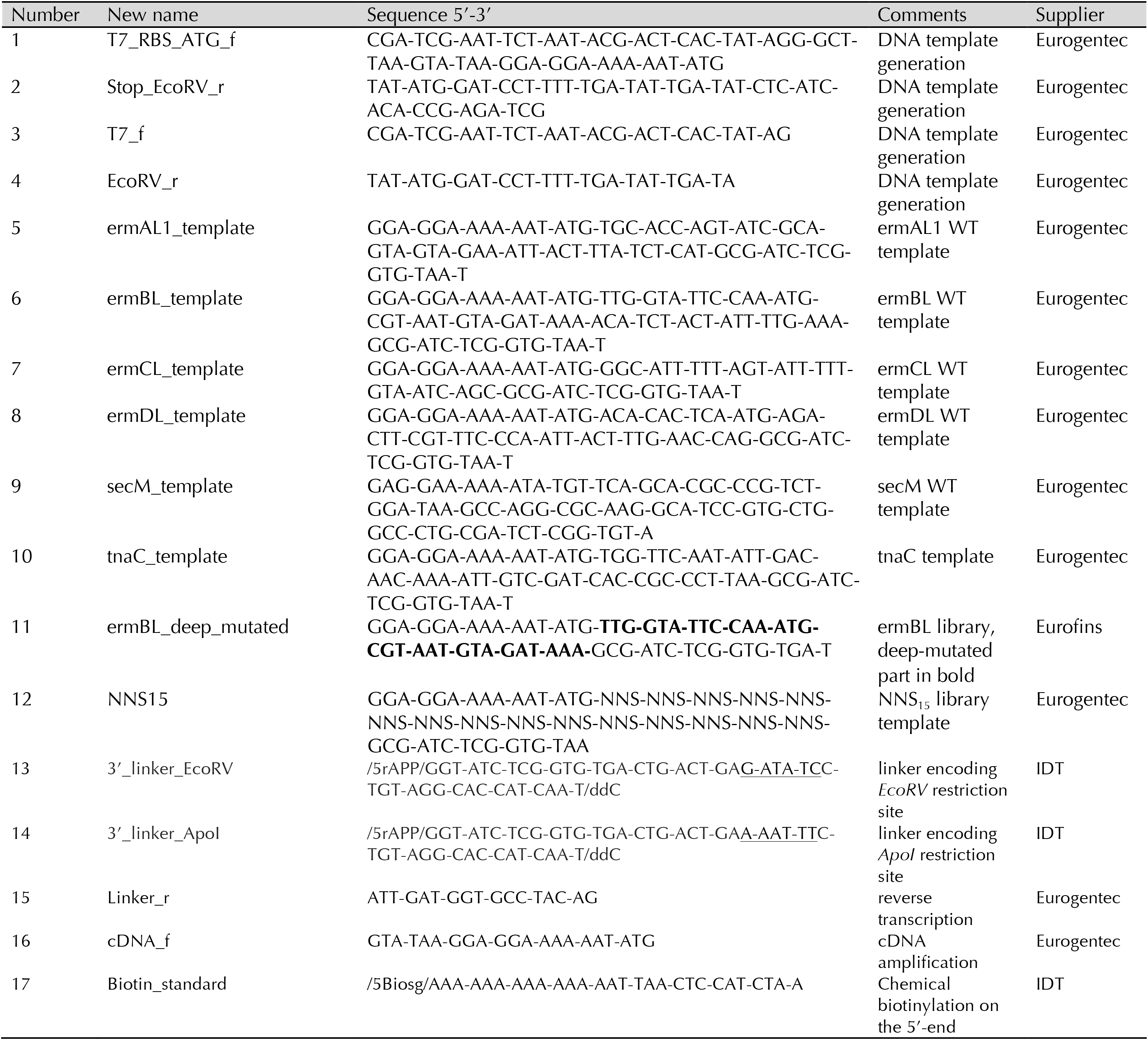
Oligonucleotides used for inverse toeprinting

**Supplementary Table 3.**
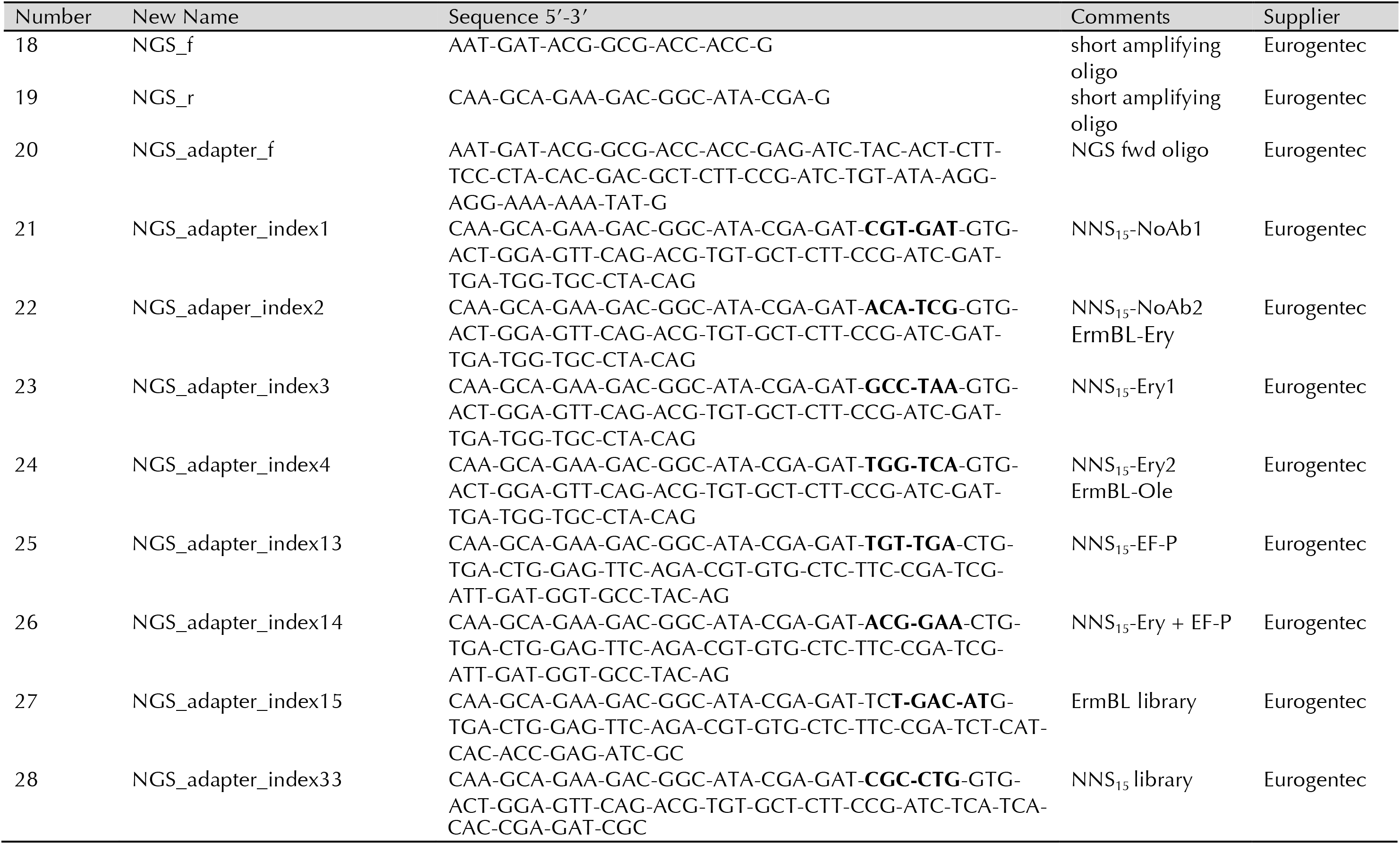
Oligonucleotides used for generating the Illumina sequencing library and barcoding

**Supplementary Table 4.**
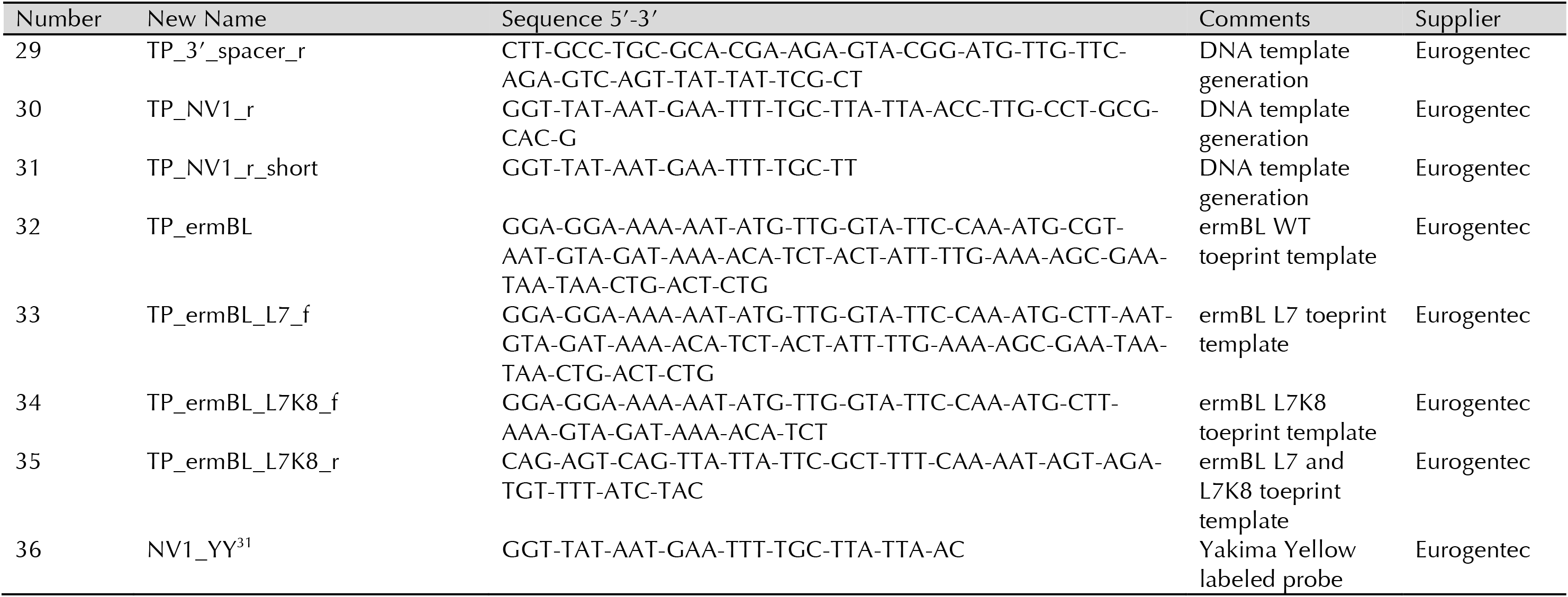
Oligonucleotides used for toeprinting

**Supplementary Table 5.**
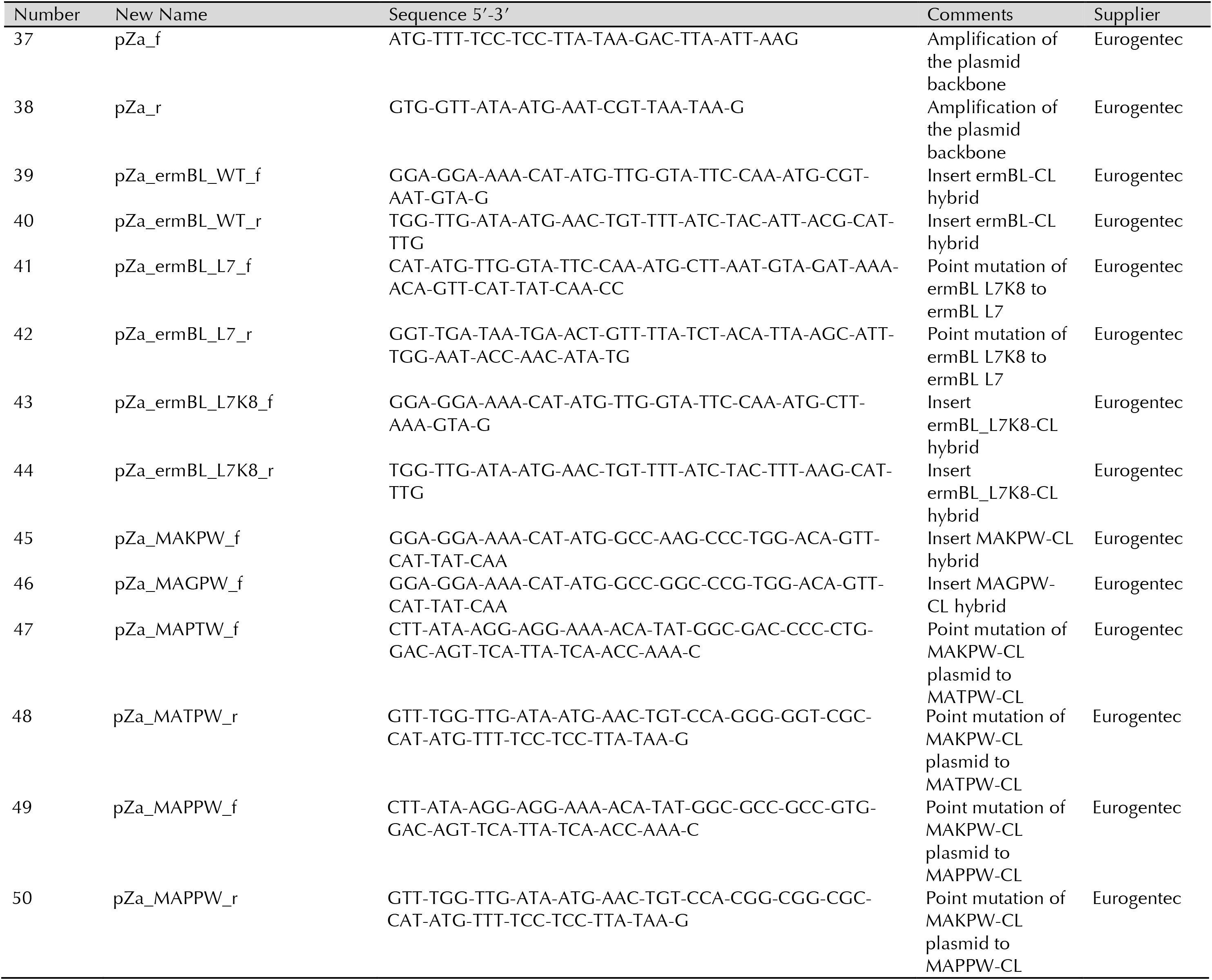
Oligonucleotides used for generating the plasmids for *in vivo* studies

**Supplementary Table 6.** Summary of NGS read processing

| Sample | Reads before filtering | Outside region of interest | Contains long 'A' stretch | First codon not ATG | Q < 30 | Reads after filtering |
| --- | --- | --- | --- | --- | --- | --- |
| NNS<sub>15</sub>-NoAb1 | 2,877,735 | 1,679,246 | 143 | 14,283 | 173,880 | 1,010,183 |
| NNS<sub>15</sub>-NoAb2 | 5,284,297 | 2,703,215 | 294 | 16,871 | 155,256 | 2,408,661 |
| NNS<sub>15</sub>-Ery1 | 4,949,363 | 2,654,430 | 162 | 15,190 | 104,646 | 2,174,935 |
| NNS<sub>15</sub>-Ery2 | 6,780,268 | 3,639,577 | 230 | 22,590 | 189,211 | 2,928,660 |
| NNS<sub>15</sub>-NoAb1<br>+EF-P | 10,765,416 | 6,024,995 | 470 | 51,040 | 790,792 | 3,898,119 |
| NNS<sub>15</sub>-Ery1<br>+EF-P | 4,515,200 | 2,062,760 | 92 | 22,025 | 411,639 | 2,018,684 |
| NNS<sub>15</sub> Library | 1,200,518 | 108,885 | 0 | 1,590 | 211,260 | 878,783 |
| ErmBL-Ery | 1,162,143 | 281,581 | 0 | 3,498 | 157,270 | 719,776 |
| ErmBL-Ole | 1,041,342 | 237,584 | 0 | 3,205 | 144,784 | 655,769 |
| ErmBL Library | 1,402,702 | 90,215 | 0 | 1,740 | 37,054 | 1,273,693 |

**Supplementary Table 7.**
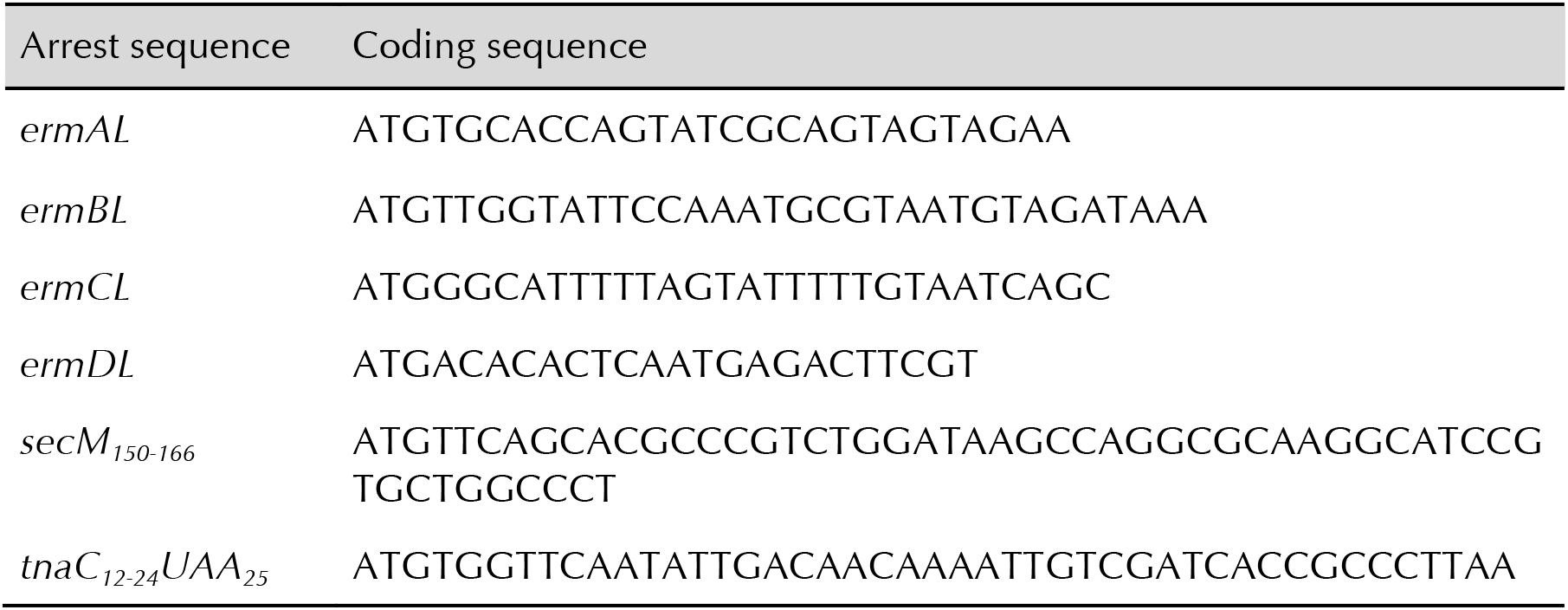
Nucleotide sequences for arrest peptides used in this study

## References

1 Kannan, K. et al. The general mode of translation inhibition by macrolide antibiotics. Proceedings of the National Academy of Sciences 111, 15958–15963 (2014).

2 Davis, A.R., Gohara, D.W. & Yap, M.-N.F. Sequence selectivity of macrolide-induced translational attenuation. Proceedings of the National Academy of Sciences of the United States of America 111, 15379–15384 (2014).

3 Marks, J. et al. Context-specific inhibition of translation by ribosomal antibiotics targeting the peptidyl transferase center. Proceedings of the National Academy of Sciences 113, 12150–12155 (2016).

4 Ramu, H., Mankin, A. & Vazquez-Laslop, N. Programmed drug-dependent ribosome stalling. Molecular Microbiology 71, 811–824 (2009).

5 Vázquez-Laslop, N., Ramu, H., Klepacki, D., Kannan, K. & Mankin, A.S. The key function of a conserved and modified rRNA residue in the ribosomal response to the nascent peptide. The EMBO Journal 29, 3108–3117 (2010).

6 Arenz, S. et al. Molecular basis for erythromycin-dependent ribosome stalling during translation of the ErmBL leader peptide. Nature communications 5, 3501 (2014).

7 Dinos, G.P. The macrolide antibiotic renaissance. British Journal of Pharmacology 174, 2967–2983 (2017).

8 Seiple, I.B. et al. A platform for the discovery of new macrolide antibiotics. Nature 533, 338 (2016).

9 Ingolia, N.T., Ghaemmaghami, S., Newman, J.R.S. & Weissman, J.S. Genome-Wide Analysis in Vivo of Translation with Nucleotide Resolution Using Ribosome Profiling. Science 324, 218–223 (2009).

10 Woolstenhulme, C.J., Guydosh, N.R., Green, R. & Buskirk, A.R. High-Precision Analysis of Translational Pausing by Ribosome Profiling in Bacteria Lacking EFP. Cell Reports 11, 13–21 (2015).

11 Hartz, D., McPheeters, D.S. & Gold, L. Selection of the initiator tRNA by *Escherichia coli* initiation factors. Genes & Development 3, 1899–1912 (1989).

12 Orelle, C. et al. Identifying the targets of aminoacyl-tRNA synthetase inhibitors by primer extension inhibition. Nucleic Acids Research 41, e144–e144 (2013).

13 Vincent, H.A. & Deutscher, M.P. Substrate recognition and catalysis by the exoribonuclease RNase R. Journal of Biological Chemistry 281, 29769–29775 (2006).

14 Shimizu, Y. et al. Cell-free translation reconstituted with purified components. Nature Biotechnology 19, 751 (2001).

15 Scolnick, E., Tompkins, R., Caskey, T. & Nirenberg, M. Release factors differing in specificity for terminator codons. Proceedings of the National Academy of Sciences 61, 768–774 (1968).

16 Horinouchi, S. & Weisblum, B. Posttranscriptional modification of mRNA conformation: Mechanism that regulates erythromycin-induced resistance. Proceedings of the National Academy of Sciences 77, 7079–7083 (1980).

17 Min, Y.-H., Kwon, A.-R., Yoon, E.-J., Shim, M.-J. & Choi, E.-C. Translational Attenuation and mRNA Stabilization as Mechanisms of *erm(B*) Induction by Erythromycin. Antimicrobial Agents and Chemotherapy 52, 1782–1789 (2008).

18 Gupta, P. et al. Nascent peptide assists the ribosome in recognizing chemically distinct small molecules. Nature chemical biology 12, 153 (2016).

19 Arenz, S. et al. A combined cryo-EM and molecular dynamics approach reveals the mechanism of ErmBL-mediated translation arrest. Nature communications 7 (2016).

20 Tuerk, C. & Gold, L. Systematic evolution of ligands by exponential enrichment: RNA ligands to bacteriophage T4 DNA polymerase. Science 249, 505–510 (1990).

21 Ellington, A.D. & Szostak, J.W. *In vitro* selection of RNA molecules that bind specific ligands. nature 346, 818 (1990).

22 Ramu, H. et al. Nascent peptide in the ribosome exit tunnel affects functional properties of the A-site of the peptidyl transferase center. Molecular cell 41, 321–330 (2011).

23 Sothiselvam, S. et al. Macrolide antibiotics allosterically predispose the ribosome for translation arrest. Proceedings of the National Academy of Sciences 111, 9804–9809 (2014).

24 Doerfel, L.K. et al. EF-P Is Essential for Rapid Synthesis of Proteins Containing Consecutive Proline Residues. Science 339, 85–88 (2013).

25 Peil, L. et al. Distinct XPPX sequence motifs induce ribosome stalling, which is rescued by the translation elongation factor EF-P. Proceedings of the National Academy of Sciences of the United States of America 110, 15265–15270 (2013).

26 Bailey, M., Chettiath, T. & Mankin, A.S. Induction of *erm(C*) expression by noninducing antibiotics. Antimicrobial agents and chemotherapy 52, 866–874 (2008).

27 Gong, F., Ito, K., Nakamura, Y. & Yanofsky, C. The mechanism of tryptophan induction of tryptophanase operon expression: Tryptophan inhibits release factor-mediated cleavage of TnaC-peptidyl-tRNAPro. Proceedings of the National Academy of Sciences 98, 8997–9001 (2001).

28 Luo, Z. & Sachs, M.S. Role of an upstream open reading frame in mediating arginine-specific translational control in *Neurospora crassa*. Journal of Bacteriology 178, 2172–2177 (1996).

29 Suzuki, H. et al. Characterization of RNase R-digested cellular RNA source that consists of lariat and circular RNAs from pre-mRNA splicing. Nucleic acids research 34, e63–63 (2006).

30 Ingolia, N.T., Brar, G.A., Rouskin, S., McGeachy, A.M. & Weissman, J.S. The ribosome profiling strategy for monitoring translation *in vivo* by deep sequencing of ribosome-protected mRNA fragments. Nature protocols 7, 1534 (2012).

31 Vazquez-Laslop, N., Thum, C. & Mankin, A.S. Molecular Mechanism of Drug-Dependent Ribosome Stalling. Molecular Cell 30, 190–202 (2008).

32 Ude, S. et al. Translation elongation factor EF-P alleviates ribosome stalling at polyproline stretches. Science 339, 82–85 (2013).

33 Cock, P.J. et al. Biopython: freely available Python tools for computational molecular biology and bioinformatics. Bioinformatics 25, 1422–1423 (2009).

34 Zhang, J., Kobert, K., Flouri, T. & Stamatakis, A. PEAR: a fast and accurate Illumina Paired-End reAd mergeR. Bioinformatics 30, 614–620 (2014).

35 Crooks, G.E., Hon, G., Chandonia, J.-M. & Brenner, S.E. WebLogo: a sequence logo generator. Genome research 14, 1188–1190 (2004).

36 Schneider, T.D. & Stephens, R.M. Sequence logos: a new way to display consensus sequences. Nucleic acids research 18, 6097–6100 (1990).

37 O’Shea, J.P. et al. pLogo: a probabilistic approach to visualizing sequence motifs. Nature Methods 10, 1211 (2013).

